# Population-level super-pangenome reveals genome evolution and empowers precision breeding in watermelon

**DOI:** 10.1101/2025.07.25.666869

**Authors:** Honghe Sun, Jie Zhang, Shengjin Liao, Shaogui Guo, Zhe Zhou, Xuebo Zhao, Shan Wu, Jiantao Zhao, Guoyi Gong, Jinfang Wang, Maoying Li, Yongtao Yu, Yi Ren, Shouwei Tian, Shaofang Li, Haiying Zhang, Sue A. Hammar, Cecilia McGregor, Robert Jarret, Patrick Wechter, Sandra E. Branham, Chandrasekar Kousik, Amnon Levi, Rebecca Grumet, Zhangjun Fei, Yong Xu

## Abstract

Pangenomes are increasingly critical for harnessing crop genetic diversity, yet their resolution and utility are often limited by insufficient sampling of high-quality genome assemblies. Here, we report a population-level watermelon super-pangenome constructed from 138 reference-grade assemblies, including 135 newly generated near-gapless genomes representing all seven watermelon species. The super-pangenome captures approximately one million structural variants (SVs), enabling accurate variant genotyping across ~900 watermelon accessions and substantially expanding variant discovery both across and within species. Broader sampling within the pangenome provides insights into genome evolution among watermelon species and sheds light on the origin of cultivated watermelon. SV-inclusive genome-wide association studies enhance trait mapping resolution and identify a copy number variation upstream of *ClFCI1* that regulates flesh color intensity in a dosage-dependent manner. Leveraging this comprehensive variation map, we developed high-accuracy genomic prediction models for 18 agronomic traits. Together, our findings and genomic resources establish a foundational framework for dissecting complex traits and accelerating precision breeding in watermelon, while offering a valuable model for SV-resolved pangenomics in crop species.

## Introduction

Watermelon (*Citrullus lanatus* subsp. *vulgaris*) is one of the most commercially important fruit crops worldwide, with a global production of approximately 105 million tonnes in 2023, ranking third among all fruit crops (FAOSTAT, 2023). Its sweet, juicy, and brightly colored flesh has driven strong consumer demand. Watermelon belongs to the genus *Citrullus*, which originates in Africa and comprises six additional extant species: *C. naudinianus*, *C. colocynthis*, *C. rehmii*, *C. ecirrhosus*, *C. amarus*, and *C. mucosospermus*. Archaeobotanical and genomic evidence indicates that sweet watermelon was domesticated in the northeastern African region from the wild form *C. lanatus* subsp. *cordophanus*^1–3^, while a recent comparative genomic analysis suggests that *C. mucosospermus* could be an additional ancestor of sweet watermelon^4^.

Modern watermelon cultivars exhibit substantial diversity in traits such as flesh color, sugar content, fruit size, shape, and rind pattern. However, intensive selection for these fruit quality attributes has led to a marked reduction in genetic diversity, accompanied by the loss of numerous disease-resistance and abiotic stress-tolerance traits, raising concerns about the long-term sustainability of watermelon production^5^. Wild *Citrullus* species possess valuable adaptive traits, including resistance to various pathogens, tolerance to environmental stresses, as well as enhanced levels of health-promoting compounds such as citrulline, which are critical resources for the genetic improvement of cultivated watermelon. Therefore, comprehensive characterization of genetic variation, both within cultivated germplasm and between cultivated and wild relatives, is urgently needed to enable effective genomics-assisted breeding and genetic engineering aimed at improving fruit quality and resilience in watermelon.

High-quality reference genomes and SNP-based variation maps have substantially advanced our understanding of watermelon domestication and key agronomic traits^6–8^. However, genomes from one or a few accessions do not offer sufficient coverage to efficiently assess the genetic diversity within a species or genus. Genetic bottlenecks in domestication and breeding have further constrained genetic diversity within cultivated watermelon germplasms. These limitations significantly restrict the genetic information available for watermelon breeding and impede the detection of causative variants/genes underlying important agronomic traits. To overcome these challenges, pangenomic approaches are essential for expanding the repertoire of genetic diversity accessible for watermelon improvement. Additionally, graph-based pangenomes greatly enhance the detection of structural variants (SVs)—including large insertions, deletions, inversions, and translocations—which comprise a substantial portion of genomic diversity and are increasingly recognized as major contributors to phenotypic variation^9–11^.

In this study, we assembled 135 near-gapless reference-quality genomes encompassing cultivated watermelon and its wild relatives. Leveraging these assemblies, we constructed a population-level graph-based super-pangenome of watermelon that captures nearly one million large SVs, which were confidently genotyped across more than 900 watermelon accessions. This resource enabled high-resolution analyses of genomic variation, providing new insights into the origin of cultivated watermelon. By integrating SVs into genome-wide association studies (GWAS) of 18 agronomic traits, we identified a causal copy number variation (CNV) underlying flesh color intensity. Furthermore, leveraging the extensive variation map within our graph-based super-pangenome, we demonstrated the potential of genomic selection for enhancing fruit quality and disease resistance in watermelon breeding. This study establishes a foundational genomic resource to accelerate future genomics-assisted breeding strategies aimed at optimizing watermelon quality and productivity.

## Results

### Genome assembly and annotation of 135 watermelon accessions

To capture the genetic diversity within the *Citrullus* genus, we selected a total of 135 representative accessions spanning all seven extant species for genome assembly. These included 1 *C. naudinianus*, 1 *C. rehmii*, 2 *C. ecirrhosus*, 5 *C. colocynthis*, 16 *C. amarus*, 9 *C. mucosospermus*, 6 *C. lanatus* subsp. *cordophanus*, and 95 *C. lanatus* subsp. *vulgaris* (comprising 7 landraces and 88 cultivars) (**Supplementary Table 1**). The panel was designed to integrate core germplasm, founder inbred lines and accessions with valuable traits such as disease resistance^12,13^. HiFi reads were generated for all 135 accessions at an average depth of ~30.3×. Additionally, Oxford Nanopore Technologies (ONT) ultra-long reads (~54.6× on average) and high-throughput chromosome conformation capture (Hi-C) reads (~153.8×) were generated for ten of these accessions, spanning cultivar, landrace, *C. lanatus* subsp. *cordophanus*, and three wild relatives widely used in disease-resistance breeding, *C. mucosospermus*, *C. amarus*, and *C. colocynthis* (**Supplementary Table 1**).

The assembled genomes had an average size of 374 Mb, with an average contig N50 size of 31.2 Mb (**Supplementary Table 1**). Approximately 99.2% of the assembled sequences were anchored and ordered onto the 11 watermelon chromosomes. Of the assembled chromosomes, 78.2% (1162 out of 1485) contained telomeres at both ends and 52.6% (781) were completely gapless. Among the ten genomes assembled from HiFi, ONT, and Hi-C reads, seven were gapless telomere-to-telomere (T2T) genomes. The remaining three contained gaps–two with a single gap and one with three–likely due to unresolved centromeric and ribosomal DNA (rDNA) repeats^14^. Genome assemblies derived from the accession ‘97103’ have served as the primary references for watermelon genomic studies. Compared to the previous version^6^, the gapless T2T genome of ‘97103’ assembled in this study (version 3) increased the size from 360 Mb to 370 Mb with markedly improved base-level accuracy (consensus quality value [QV] of 65 vs. 45.3).

BUSCO^15^ evaluation revealed high completeness across these 135 genome assemblies, with an average completeness rate of 99.1%. Assessment using a k-mer-based approach^16^ indicated an average QV of 64.9 (**Supplementary Table 1**). These metrics were comparable to or higher than those reported for the recently published watermelon genome assemblies^4^, underscoring the robustness and high accuracy of our assemblies. The transposable element (TE) content ranged from 59.0% to 64.7%, and between 20,834 and 23,330 protein-coding genes were predicted in these 135 *Citrullus* genomes (**Supplementary Table 2**).

### Chromosomal evolution of the *Citrullus* genus

Large chromosomal rearrangements, including translocations and inversions, play crucial roles in plant evolution and domestication^17,18^. Understanding these structural variations in *Citrullus* accessions can enhance their effective utilization in breeding programs. Comparative analysis of the 135 genome assemblies generated in this study, along with three previously published genomes^2,19,20^, identified a total of 11 translocations and 101 large inversions (>100 kb) (**Fig. 1a** and **Supplementary Tables 3 and 4**). Five of the translocations were specific to *C. lanatus*. Notably, one event between chromosomes 2 and 3 has been reported to disrupt the structure of the gynoecious gene *ClWIP1*, thereby leading to the gynoecious phenotype^21^. Another *C. lanatus*-specific translocation, between chromosomes 6 and 10, has been found to cause chromosomal synapsis abnormalities during meiotic diakinesis in hybrids, resulting in fruits with reduced seed numbers^22^. Of the 101 large inversions, 52 were specific to wild relatives (*C. colocynthis*, *C. amarus*, and *C. mucosospermus*), with 22 overlapping with known disease-resistance QTLs (**Supplementary Table 4**).

**Figure 1.**
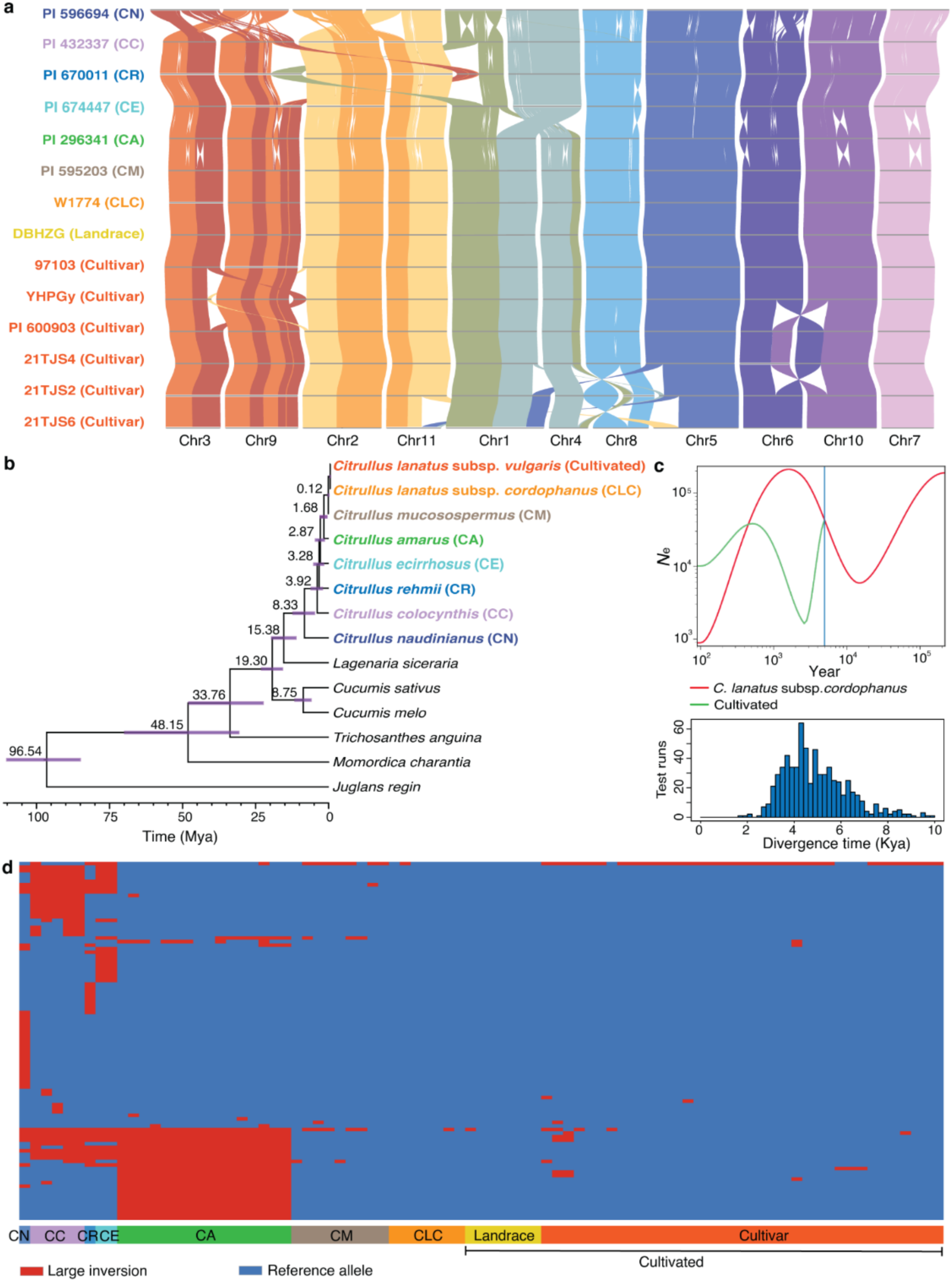
Genome evolution in the *Citrullus* genus. (**a**) Whole-genome alignments showing inter-chromosomal translocations and large inversions. One representative genome from each wild species/subspecies (abbreviations defined in **b**), along with genomes from one landrace and five cultivars, are displayed. (**b**) Time-calibrated species tree. (**c**) Estimated divergence time between *C. lanatus* subsp. *cordophanus* and cultivated watermelon using SMC++. Top: Demographic history of the effective population size in *C. lanatus* subsp. *cordophanus* and cultivated watermelon. The blue vertical line marks the estimated split time between the two. Bottom: Histogram of split time estimates based on random accession sampling, with a mean divergence time of ~4.9 thousand years ago (Kya). (**d**) Distribution of large inversions across genome assemblies (53 cultivars without large inversions are not shown).

Wild relatives have long been recognized as invaluable source for disease resistance in watermelon. However, linkage drag often complicates the breeding process by inadvertently introducing undesirable alleles, leading to trade-offs in key traits such as yield and fruit quality. For instance, QTLs associated with Fusarium wilt resistance (*qFon2-6*; 9.77-25.00 Mb), sweetness (*QBrix6*; 10.21-11.24 Mb), and flesh firmness (13.00-20.54 Mb) are closely located on chromosome 6 (refs. ^23–25^). Introgression analysis using RFMix^26^ revealed that the genomic region encompassing these three QTLs originated from *C. amarus* (**Supplementary Fig. 1**). Three large inversions within this region (10.49-13.15 Mb, 17.43-19.64 Mb, and 21.96-23.29 Mb) likely suppress recombination^27^, resulting in the acquisition of desirable traits like disease resistance and firm flesh but at the cost of decreased sweetness. The cultivar ‘SugarleeXZ’ inherited all these three inversions. In contrast, *C. amarus* accessions ‘PI 296341-FR’ and ‘USVL252’ lacked the first and the first two inversions, respectively (**Supplementary Fig. 1**). Therefore, using ‘PI 296341-FR’ and ‘USVL252’ in backcross breeding could facilitate the introgression of resistance and firmness traits without compromising sweetness.

The evolutionary timeline and genomic relationships among *Citrullus* species reconstructed in this study (**Fig. 1b**) broadly aligned with previous reports^4,28^. Notably, we estimated that *C. mucosospermus* diverged from *C. lanatus* approximately 120,000 years ago, well before the domestication of any crops (<12,000 years ago)^29^. In contrast, the divergence between domesticated watermelon and *C. lanatus* subsp. *cordophanus* was estimated at ~4,900 years ago (**Fig. 1c**), closely aligning with archaeological evidence of watermelon domestication (~4,000– 6,000 years ago) in northeastern Africa^20,30^. These evolutionary timelines further support *C. lanatus* subsp. *cordophanus* as the likely direct wild progenitor of cultivated watermelon, while *C. mucosospermus* is unlikely to be a direct ancestor.

A recent study proposed *C. mucosospermus* as an additional progenitor based on seven diagnostic variants shared with cultivated watermelon but absent from *C. lanatus* subsp. *cordophanus*^4^. However, this conclusion relied on a single *C. lanatus* subsp. *cordophanus* genome, overlooking potential intraspecific variation within this subspecies. In this study, we analyzed seven *C. lanatus* subsp. *cordophanus* genome assemblies, revealing that four of the seven variants were, in fact, segregating within the population (**Supplementary Table 5**). ABBA-BABA tests^31^ revealed that the remaining three variants did not fall within genomic regions introgressed from *C. mucosospermus*, suggesting that they are unlikely derived from that species (**Supplementary Table 6**). Thus, all seven variants can be plausibly explained by variation within *C. lanatus* subsp. *cordophanus*, and the absence of three from current assemblies likely reflects incomplete sampling. These findings highlight the importance of comprehensive population sampling when inferring crop ancestry, as reliance on limited genomes can obscure intraspecific diversity and lead to misleading conclusions.

### Comprehensive gene-based super-pangenome

Through gene clustering across the 138 *Citrullus* genomes, we constructed a comprehensive gene-based super-pangenome for wild and cultivated watermelons, comprising 35,919 pangenes– approximately 1.6 times the number of genes in the ‘97103’ reference genome (v3). The number of pangenes increased with the inclusion of additional genomes, plateauing around 80 (**Fig. 2a**). Pangenes were classified into four categories: core (39.3%), softcore (5.4%; presented in 137 accessions), shell (54.4%; present in 2-136 accessions), and private (0.9%) (**Fig. 2b**). Within domesticated watermelon, the inclusion of 96 genomes identified 7,279 additional pangenes beyond those found in the ‘97103’ genome (**Supplementary Fig. 2**). As the wild progenitor of cultivated watermelon, *C. lanatus* subsp. *cordophanus* contributed 1,537 (4.3%) additional pangenes absent in cultivated accessions (**Fig. 2c** and **Supplementary Fig. 2a**). Three wild relatives widely used in watermelon breeding programs, *C. mucosospermus*, *C. amarus*, and *C. colocynthis*, collectively contributed 4,732 pangenes (13.2%) not found in *C. lanatus*. In contrast, *C. naudinianus*, *C. rehmii*, and *C. ecirrhosus* contributed only 666 additional pangenes, likely due to limited sampling.

**Figure 2.**
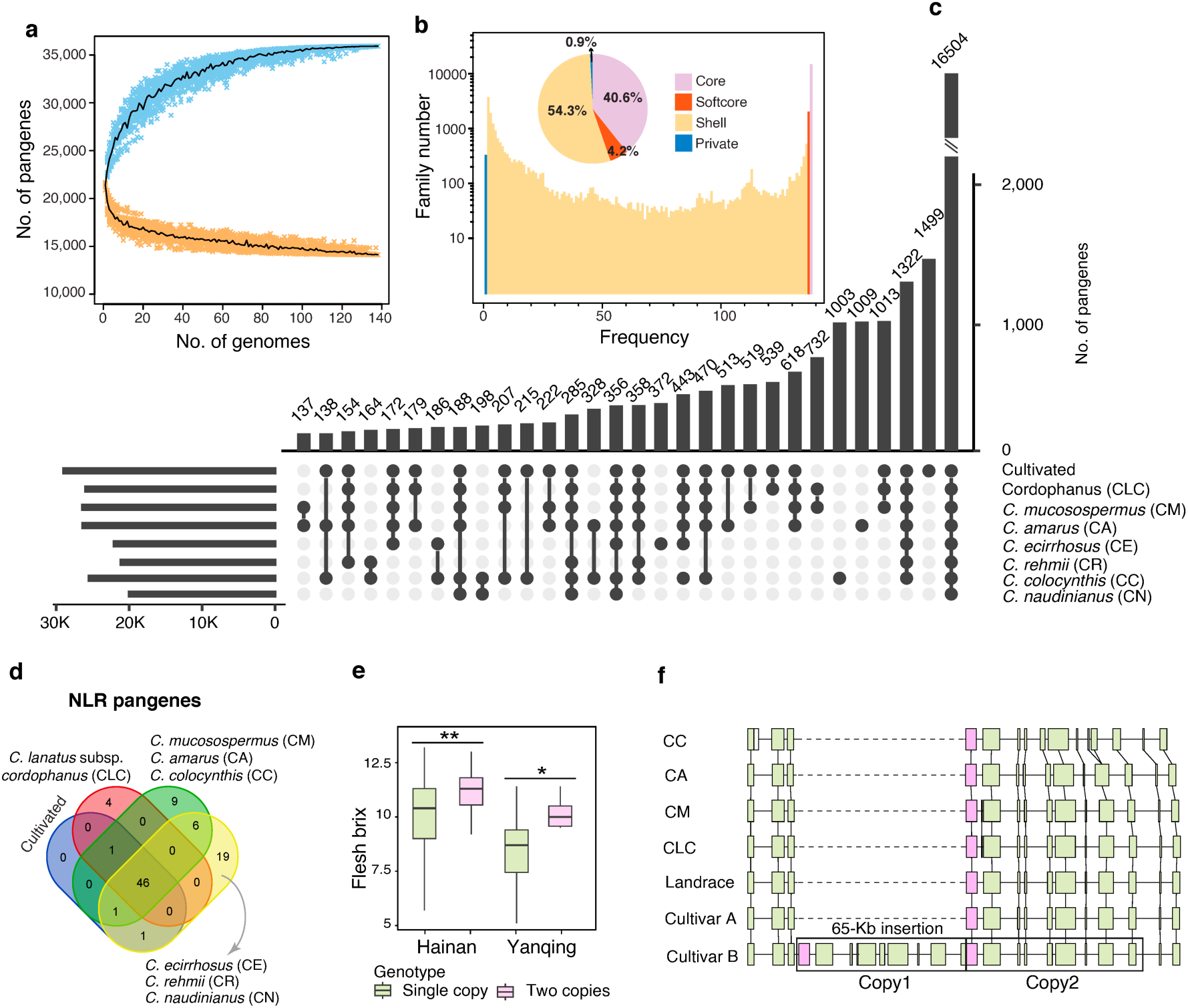
Gene pool dynamics in the *Citrullus* genus. (**a**) Modeling of gene-based pan- and core-genome sizes as additional genomes are incorporated. (**b**) Composition of the gene-based *Citrullus* super-pangenome. (**c**) Presence-absence variation (PAV) of pangenes across cultivated watermelon and its wild relatives. Top 30 intersected groups are plotted. (**d**) Presence-absence variation of NLR pangenes across *Citrullus* species. (**e**) Flesh sugar content (°Brix) in accessions carrying one or two copies of the hexosyltransferase gene. Data are from two field trials, conducted in Hainan and Yanqing. ‘*’ and ‘**’ indicate *P* < 0.05 and *P* < 0.01, respectively (Student’s *t* test). (**f**) Schematic diagram of the cultivar-specific 65-kb insertion in different *Citrullus* groups. Each rectangle represents a gene, and purple rectangles indicate the hexosyltransferase gene. Cultivar A: ‘97103’; Cultivar B: ‘TWFYingRou’.

Nucleotide-binding site leucine-rich repeat (NLR) genes play a pivotal role in plant disease resistance^32^. In addition to the 46 NLR pangenes present in the ‘97103’ reference genome, we identified 41 novel NLR pangenes from the pangenome: 3 from non-reference cultivated watermelons, 4 from *C. lanatus* subsp. *cordophanus*, and 34 from six wild species (**Fig. 2d** and **Supplementary Table 7**), highlighting wild species as a rich reservoir of disease resistance diversity.

Tandem gene duplication can increase gene dosage and enhance the expression of beneficial traits. Using the gene-based super-pangenome, we identified 31 pangenes that exhibited higher frequencies of tandem duplication in cultivated watermelons compared to wild species and were expressed at higher levels in the flesh of ‘97103’ relative to the wild accession ‘PI 296341-FR’ (**Supplementary Table 8**). These included the previously reported tandem duplication of *ClTST2*, which is associated with increased flesh sweetness and was strongly favored during domestication, becoming nearly fixed in cultivars (allele frequency of 97.1%)^2^, thereby limiting its potential in future improvement of fruit sweetness. Among the newly identified tandem duplicates, we discovered a hexosyltransferase gene duplication (*XG0025C01G000760* and *XG0025C01G000860*) specific to cultivars, with an allele frequency of 28%. Hexosyltransferases catalyze the transfer of hexose sugars to various acceptor molecules and are involved in sucrose metabolism^33^. Notably, we found that this duplication was significantly associated with increased flesh sweetness (**Fig. 2e**). These findings suggest that the hexosyltransferase duplication represents a promising target for enhancing flesh sweetness in future watermelon breeding.

### Graph-based pangenome facilitates trait-variation association

To capture the full spectrum of genetic diversity within the *Citrullus* genus, we performed pairwise genome alignments using the ‘97103’ genome as the reference, leading to the identification of 37,699,340 SNPs, 8,294,544 small indels (<20 bp), and 910,844 large SVs (≥20 bp; including 502,800 insertions and 408,044 deletions). The number of SVs per accession was positively correlated with their genetic distance from ‘97103’ (**Fig. 3a** and **Supplementary Table 9**). The cumulative SV count within each group significantly surpassed the average observed in individual accessions (**Fig. 3b**). Notably, the inclusion of 89 cultivars captured 60,297 SVs–over ten times the average (5,933)–highlighting a substantially enhanced representation of genetic diversity.

**Figure 3.**
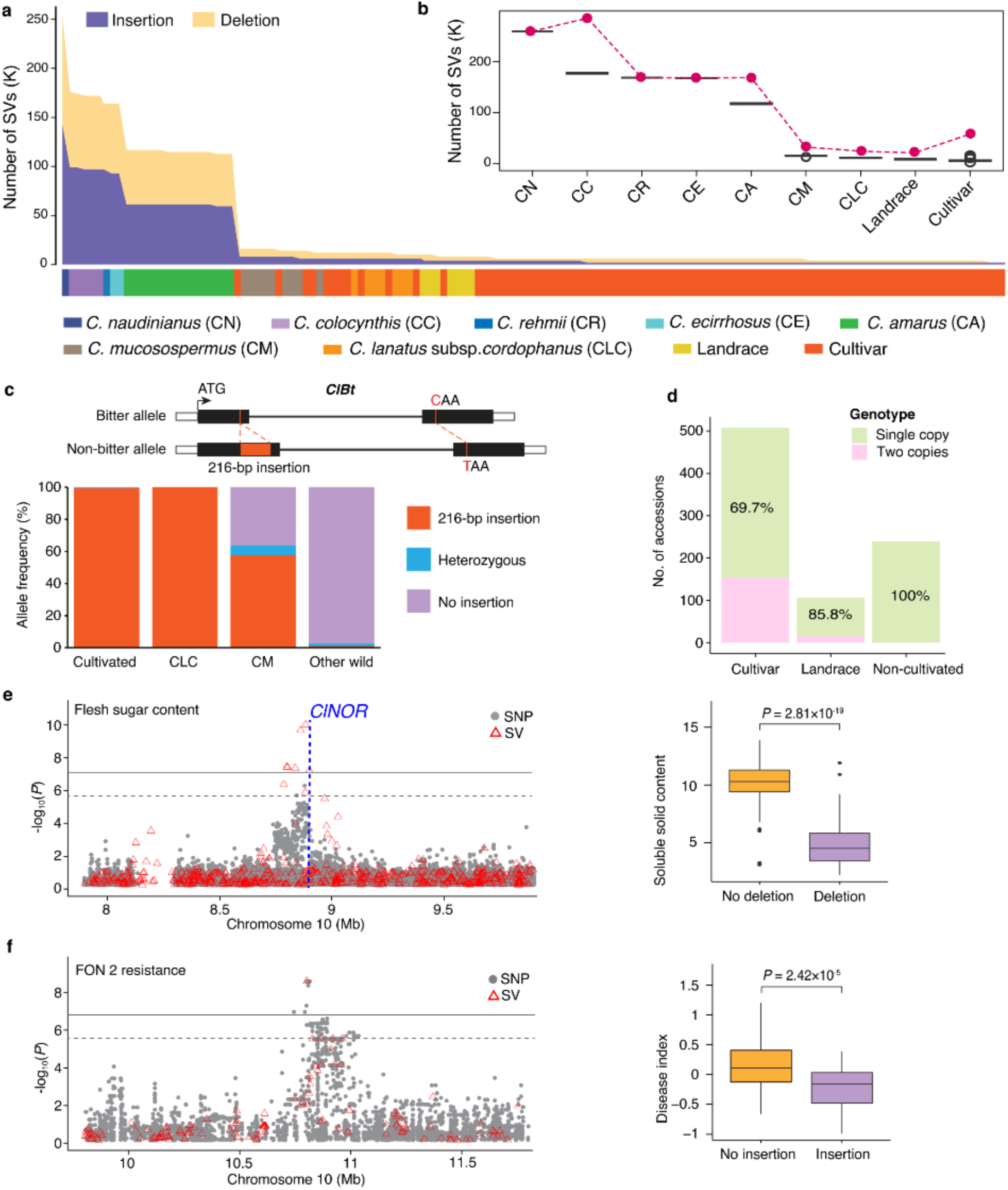
Landscape of structural variation across *Citrullus* species. (**a**) Number of large insertions and deletions identified in each of the 138 *Citrullus* accessions. (**b**) Average (black line) and cumulative (red dots) number of distinct structural variants (SVs) across different *Citrullus* groups. Detailed numbers are provided in **Supplementary Table 9**. (**c**) Haplotypes of the *ClBt* gene and their distribution among different watermelon groups. (**d**) Copy number variation of the 65-kb insertion across cultivars, landraces, and wild *Citrullus* accessions. (**e,f**) Local Manhattan plots (left) and corresponding box plots showing the distribution of accessions carrying distinct alleles (right) for flesh sugar content (**e**) and FON 2 (*Fusarium oxysporum* f. sp. *niveum* race 2) resistance (**f**). Horizontal solid and dashed lines represent Bonferroni-corrected genome-wide significance thresholds at α = 0.05 and α = 0.10, respectively.

We then constructed a graph-based pangenome by integrating SNPs, indels, and SVs, enabling accurate SV genotyping in 776 re-sequenced accessions, including 313 newly sequenced in this study (**Supplementary Table 10**). Combined with variants from the 138 genome assemblies, this yielded a comprehensive variation map encompassing 914 wild and cultivated watermelon accessions. Using this SV-inclusive variation map, we uncovered a previously unreported 216-bp insertion in the first exon of *ClBt* (the bitterness gene), which introduces three premature stop codons. This insertion was in complete linkage with the previously reported nonsense SNP in the second exon^6^, together forming a haplotype that underlies the loss of bitterness. This haplotype was completely fixed in cultivated watermelons and *C. lanatus* subsp. *cordophanus*, but was rare in wild relatives, particularly those other than *C. mucosospermus* (**Fig. 3c**). The discovery of this insertion provides new insight into the genetic basis of bitterness loss and reveals a key haplotype at the *ClBt* locus.

We further identified 622 and 983 SVs under selection during watermelon domestication (*C. lanatus* subsp. *cordophanus* vs. landrace) and improvement (landrace vs. cultivar), respectively, encompassing 184 and 395 genes. Among SVs under selection during improvement, we identified a 65-kb insertion that led to the tandem duplication of the aforementioned hexosyltransferase gene (**Fig. 2f**). Allele frequency analysis indicated that this insertion originated in landrace and was subsequently favored during watermelon improvement (**Fig. 3d**). Additionally, this variation map provided insights into the genetic basis of fruit shape: both the nonsynonymous SNP and 159-bp deletion in *ClFS1* (a key gene for fruit shape) reported previously^34,35^ were captured. The nonsynonymous SNP was present in both *C. amarus* (allele frequency of 42%) and cultivated watermelons (11%), while the 159-bp deletion was found exclusively in cultivated watermelons (9%), indicating that this variation likely originated during domestication (**Supplementary Fig. 3a,b**). Furthermore, the phenotypic effect of the deletion appeared to be stronger than that of the SNP (**Supplementary Fig. 3c**). Together, these findings highlight how the graph-based pangenome elucidates key domestication and improvement alleles underlying important traits such as fruit bitterness, sweetness, and morphology in watermelon.

Empowered by the comprehensive variant set–particularly SVs–captured in the graph-based pangenome, we conducted genome-wide association studies (GWAS) for 12 fruit-quality and 6 disease-resistance traits (**Supplementary Table 11** and **Supplementary Figs. 4–5**). We identified a total of 93 loci significantly associated with at least one trait, 14 (14.9%) of which had lead signals marked by SVs (**Supplementary Table 12**). Notably, a GWAS signal for flesh sweetness on chromosome 10 was detected exclusively in the SV-based GWAS. This signal included a 29-bp deletion located 187 bp downstream of the transcription factor *ClNOR* (*XG0025C10G005980*) (**Fig. 3e**), a gene recently shown to regulate fruit ripening and sugar accumulation in watermelon^36^. For resistance to *Fusarium oxysporum* f. sp. *niveum* (FON) race 2, a 135-bp insertion at 10.8 Mb on chromosome 10 showed the strongest association, and accessions carrying this insertion exhibited significantly enhanced resistance (**Fig. 3f**). This SV is located 4,224 bp upstream of an AAA-type ATPase gene (*XG0025C10G006750*), whose homologs have been implicated in broad-spectrum disease resistance in rice and *Arabidopsis*^37,38^. For nematode resistance, an association signal was detected on chromosome 3, overlapping with the previously mapped QTL 3.1 for nematode resistance^39^. The lead variant was an SV located 7,014 bp upstream of *XG0025C03G016370*, which encodes a calmodulin-binding protein 60 (CBP60), a member of a tandem gene cluster in this region. Additional associated SVs were identified within the promoter, intronic, and coding regions of other *CBP60* members in the cluster (**Supplementary Fig. 6**). Given that *CBP60* genes are known regulators of plant immune responses^40^, this locus likely plays a pivotal role in nematode defense. Together, these results underscore the enhanced resolution and discovery power of SV-inclusive GWAS in uncovering trait-associated variants in watermelon.

### A copy number variant regulates flesh color intensity

Flesh color is a key fruit-quality trait in watermelon, with brighter flesh colors enhancing visual appeal and reflecting higher levels of nutritional compounds such as carotenoids. Using chroma value as a quantitative metric (**Fig. 4a**), our GWAS identified two significant signals for flesh color intensity: a major locus on chromosome 6 at 24.5 Mb (*FCI1*) and a minor one on chromosome 8 at 24.9 Mb (*FCI2*) (**Fig. 4b**). The major peak at *FCI1* corresponded to a 2,516-bp insertion that exhibited low linkage disequilibrium with nearby SNPs, likely explaining why it was not detected in SNP-based GWAS (**Fig. 4c**). Bulked-segregant analysis (BSA) using an F2 population derived from a cross between ‘Ming 58’ (scarlet red flesh) and ‘JX2’ (pink flesh) independently mapped the *FCI1* locus to a 2.8-Mb interval (23.8–26.6 Mb) (**Supplementary Fig. 7a**). Fine mapping using this F_2_ population (n = 141) and an additional F_2_ population derived from ‘JLM’ (yellow flesh) × ‘Cream of Saskatchewan’ (pale yellow flesh) (n = 636) narrowed the locus to a 146-kb region (24.45-24.60 Mb) containing 13 genes (**Supplementary Fig. 7b,c**). Of these, five were expressed in fruit flesh and only *XG0025C06G012030* (hereafter *ClFCI1*) showed an expression pattern consistent with contrasting dark- and light-flesh phenotypes (**Supplementary Fig. 7d**). *ClFCI1*, which encodes a tetratricopeptide repeat (TPR) protein, is homologous to the Arabidopsis REDUCED CHLOROPLAST COVERAGE (REC) genes known to regulate chloroplast compartment size and chlorophyll content^41^. The 2,516-bp insertion corresponding to the *FCI1* peak resulted in a tandem triplication of a 1,258-bp promoter segment located ~1.8 kb upstream of *ClFCI1* (**Fig. 4d**). It is worth noting that no other sequence polymorphisms were found within or near the *ClFCI1*coding region between the mapping parents.

**Figure 4.**
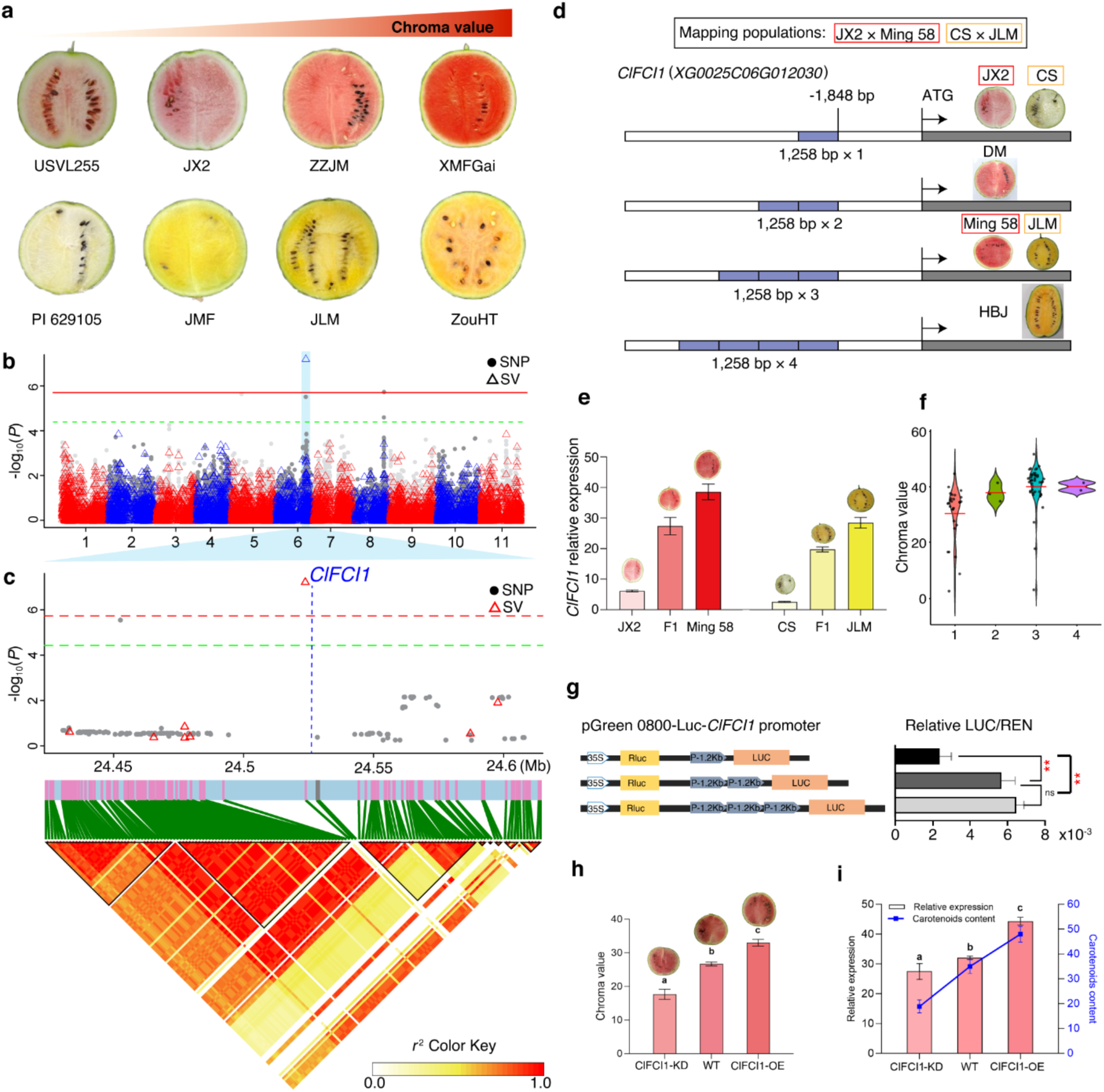
Copy number variation in the promoter region of *ClFCI1* controls watermelon flesh color intensity. (**a**) Photos of representative watermelon accessions showing the gradient of flesh color intensity. (**b**) Manhattan plot of GWAS for flesh color intensity. Red and green horizontal lines indicate genome-wide significance thresholds at α = 0.05 and α = 0.10, respectively. (**c**) Zoomed-in Manhattan plot (top) and LD heatmap (bottom) at the *ClFCI1* locus (24.5-24.6 Mb on chromosome 6). Black triangles in LD heatmap indicate LD blocks. The 2,516-bp insertion was the only variant in this region significantly associated with flesh color intensity. (**d**) Structural diagram showing copy-number variation of the 1,258-bp segment (blue boxes) in the *ClFCI1* promoter across representative watermelon accessions. (**e**) Expression levels of *ClFCI1* in fruit flesh of parents and F_1_ lines from the crosses ‘Ming 58’ × ‘JX2’ (left) and ‘JLM’ × ‘CS’ (‘Cream of Saskatchewan’; right). Error bars indicate the standard deviation of three biological replicates. (**f**) Violin plots of chroma values in accessions carrying one to four copies of the 1,258-bp segment. Black dots indicate chroma values in individual accessions, and horizontal red bars indicate mean chroma values. (**g**) Dual-luciferase (LUC) reporter activity driven by *ClFCI1* promoters carrying one to three copies of the 1,258-bp segment. Error bars indicate standard deviation of ten biological replicates. ** denotes *P* < 0.01 (Student’s *t*-test). (**h**) Representative fruits and chroma values of *ClFCI1*-knockdown (ClFCI1-KD), wild type (WT, ‘ZZJM’), and *ClFCI1*-overexpression (ClFCI1-OE) lines. (**i**) Carotenoid contents (mg/kg flesh weight) and relative *ClFCI1* expression in wild-type and transgenic lines. Fruits were sampled at 34 days after pollination. Error bars indicate standard deviation of three biological replicates. Different lowercase letters indicate significant differences according to Turkey’s multiple range test (*P* < 0.05).

Different copy numbers of the 1,258-bp promoter segment, ranging from one to four, were observed among watermelon accessions (**Fig. 4d**). The copy number of this segment showed a positive correlation with both flesh color intensity and *ClFCI1* transcript abundance in the mapping parents and their F_1_ hybrids (**Fig. 4e**). Similarly, across the panel of natural watermelon accessions, increased copy number was positively associated with greater flesh color intensity (**Fig. 4f**). Transient dual-luciferase assays further validated a dose-dependent increase in promoter activity, with multi-copy alleles driving significantly higher reporter expression compared to the single-copy allele (**Fig. 4g**).

To elucidate the functional role of *ClFCI1* in regulating flesh color intensity, we generated both antisense knockdown and overexpression lines. Knockdown of *ClFCI1* in the red-fleshed line ‘ZZJM’ reduced pigmentation and carotenoid content in the fruit flesh. In contrast, overexpression of *ClFCI1* enhanced flesh pigmentation and increased carotenoid accumulation (**Fig. 4h,i**). Transcriptome analysis of fruits at 34 days after pollination (DAP) identified 1,131 and 832 differentially expressed genes (DEGs) in the *ClFCI1* knockdown and overexpression lines, respectively, compared to the wild type (**Supplementary Tables 13 and 14**). These DEGs were significantly enriched for genes involved in photosynthesis and plastid development (**Supplementary Table 15**), suggesting that *ClFCI1* regulates watermelon flesh color intensity primarily by modulating these processes.

Across the 914-accession panel, multi-copy alleles of the 1,258-bp *ClFCI1*-promoter segment were predominantly found within *C. lanatus*, while in wild *Citrullus* species, only single instances of tandem duplication and triplication were observed in *C. mucosospermus* and *C. amarus*, respectively (**Supplementary Table 16**). Within *C. lanatus*, the frequency of multi-copy alleles increased significantly from 12.5% in the wild progenitor *cordophanus* to 50.25% in domesticated accessions (Fisher’s exact test, *P* = 0.0037), and showed a moderate increase during improvement, from 40.38% in landraces to 51.74% in modern cultivars (*P* = 0.14). Among cultivars, tandem triplication was more prevalent than tandem duplication, consistent with selection favoring intensified flesh coloration in modern watermelon breeding programs. These findings suggest that copy number variation in the *ClFCI1* promoter represents a promising target for marker-assisted breeding aimed at enhancing flesh color intensity and nutritional value in watermelon.

### Genomic selection empowered by the graph pangenome

Genomic selection has emerged as a transformative strategy for accelerating crop improvement^42^. Here, leveraging the comprehensive variation map captured in the graph pangenome, we established a robust genomic selection framework for watermelon by training prediction models for 18 traits related to fruit quality and disease resistance. For each trait, we employed CropGBM^43^ for both marker selection and genomic prediction, identifying informative genome-wide variants based on feature importance. We built genomic selection models using either an SNP-only panel or a combined SNP+SV panel. Most traits required 476–756 markers, with two traits needing fewer than 100 (**Supplementary Table 17**). Five-fold cross-validation with five repeats yielded generally high prediction accuracies, ranging from 0.56 to 0.97 (**Fig. 5a**). Inclusion of SVs did not improve the prediction accuracies for 17 of the 18 traits but did slightly enhance performance for the flesh color category trait, indicating that large-effect structural polymorphisms not sufficiently captured by SNPs alone may underlie this trait.

**Figure 5.**
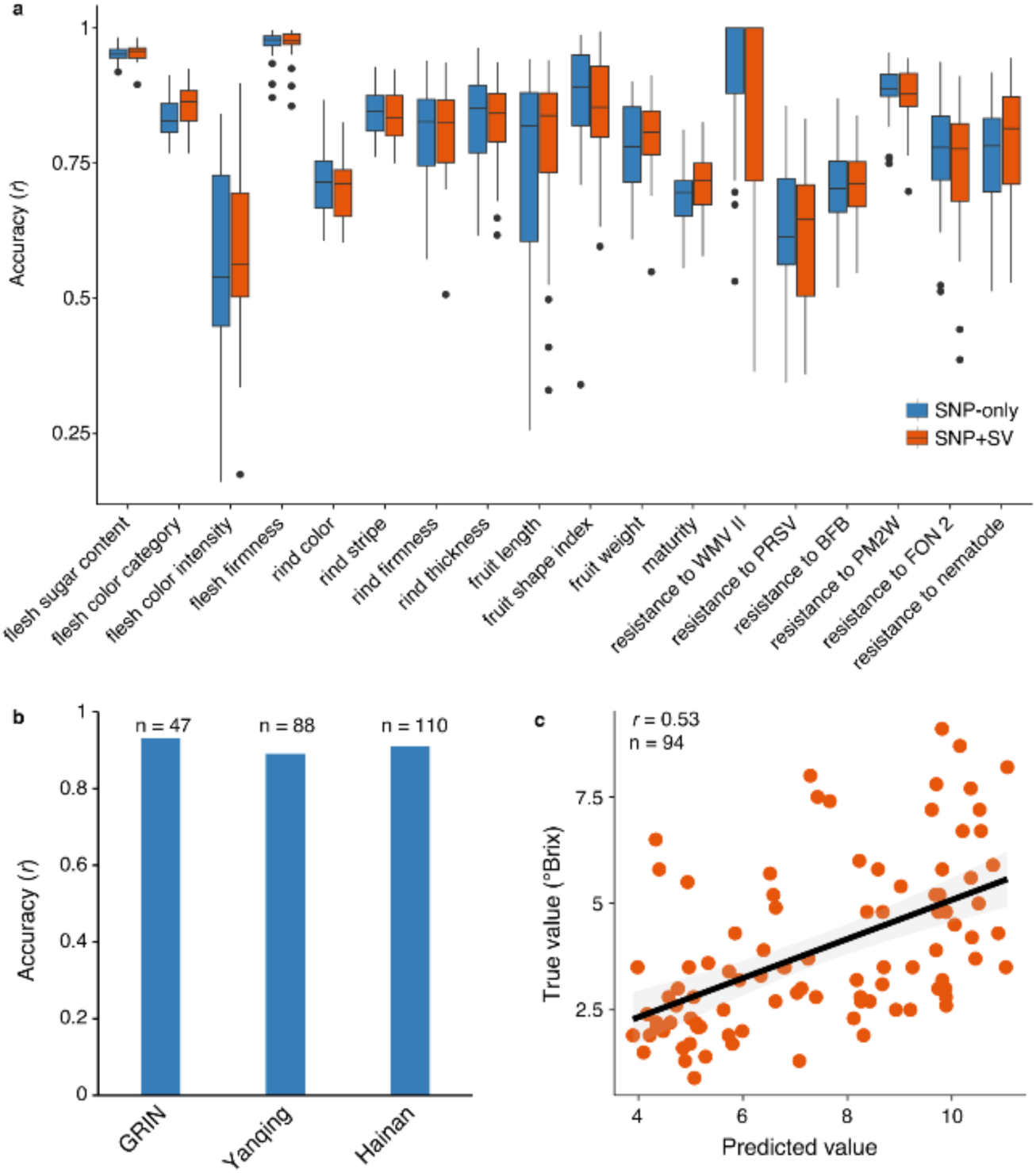
Graph-based pangenome empowers genomic selection in watermelon. (**a**) Genomic prediction accuracies for 12 fruit-quality traits and 6 disease-resistance traits using models built with high-effect variants selected from two different sets: SNP-only and SNP+SV. Trait abbreviations: WMV II, resistance to watermelon mosaic virus II. PRSV, resistance to *papaya ringspot* virus-watermelon strain. BFB, resistance to bacterial fruit blotch. PM2W, resistance to powdery mildew race 2W. FON 2, resistance to *Fusarium oxysporum* f. sp. *niveum* race 2. For each boxplot, the lower and upper bounds indicate the first and third quartiles, respectively, the center line indicates the median, and the whiskers extend to 1.5× the interquartile range. (**b**) Genomic prediction accuracies for flesh sugar content (°Brix) in cultivated watermelon using the model built with selected SNP+SV markers. Flesh sugar contents were measured in three independent experiments: GRIN, data from the USDA GRIN database; Yanqing, field trial in 2019 in Yanqing, China; Hainan; field trial in 2022 in Hainan, China. (**c**) Linear regression analysis of predicted and observed flesh sugar contents in a RIL population derived from a cross between cultivar ‘97103’ and *C. amarus* ‘PI 296341-FR’.

To further validate model robustness, we focused on the flesh sugar content trait and tested prediction accuracy using datasets of non-overlapping accessions from three phenotyping experiments independent from training: historical records from the USDA GRIN database, a 2019 field trial in Yanqing, China^6^, and a 2022 field trial in Hainan, China conducted in this study. Without retraining, the SNP+SV-based genomic selection model achieved consistently high accuracies across these datasets, ranging from 0.89 to 0.93 (**Fig. 5b**). To simulate real-world breeding applications, we further evaluated the model in a recombinant inbred line (RIL) population derived from a cross between the sweet cultivar ‘97103’ and the non-sweet *C. amarus* accession ‘PI 296341-FR’. The F_1_ hybrids, same as ‘PI 296341-FR’, exhibited the non-sweet phenotype, indicating epistatic suppression of sweetness—a challenge for genomic prediction. Despite this, the SNP+SV model maintained a moderate prediction accuracy of 0.53 (**Fig. 5c**).

Collectively, these results demonstrate that our compact marker panels (~500–700 high-effect variants) enable stable genomic predictions in watermelon. The inclusion of SVs provides limited additional predictive power, likely because most SVs are effectively linked to nearby SNPs.

## Discussion

In this study, we assembled 135 high-quality, near-gapless genomes representing all extant *Citrullus* species and constructed a population-scale graph-based super-pangenome. This enabled accurate detection and genotyping of nearly one million SVs across 914 accessions, substantially expanding upon previous genomic and pangenomic resources of watermelon^2,4,6^. The breadth and quality of these genomic resources allowed us to re-examine the origin of cultivated watermelon, providing strong evidence that *C. lanatus* subsp. *cordophanus* is the direct wild progenitor. In contrast to the findings of the previous pangenomics study^4^, our broader sampling suggests that *C. mucosospermus* is unlikely to be an additional direct ancestor of cultivated watermelon. Our findings highlight the importance of extensive intraspecific sampling for accurately reconstructing domestication history and capturing genetic diversity, providing valuable insights for leveraging wild *Citrullus* species in watermelon resistance breeding.

Flesh color is a prominent quality trait in watermelon. While previous studies have primarily focused on identifying loci that control discrete color categories^44–50^, the genetic basis of flesh color intensity—a quantitative trait relevant to both breeding objectives and consumer preferences—has remained poorly understood. Here, leveraging accurate SV genotyping enabled by the graph-based super-pangenome, we conducted GWAS and identified a copy number variant (CNV) involving a 1,258-bp sequence in the promoter of *ClFCI1* that modulates flesh color intensity. This CNV, present in one to four tandem copies, was strongly associated with *ClFCI1* expression and, consequently, with flesh color intensity and carotenoid accumulation. Notably, the frequency of multi-copy alleles increased during watermelon domestication and improvement, consistent with breeding preferences for brighter flesh colors. Transcriptomic analyses of *ClFCI1* knockdown and overexpression lines revealed that *ClFCI1* regulates flesh color intensity primarily by modulating genes involved in photosynthesis and plastid development. These findings identified a novel regulatory variant of both functional and breeding relevance and demonstrate the power of SV-informed pangenomics in uncovering the genetic basis of complex trait variation.

The comprehensive variation map also enabled genomic prediction across a broad range of fruit-quality and disease-resistance traits. For most traits, compact marker panels composed of several hundred high-effect variants achieved high predictive accuracy. Notably, the model for flesh sugar content performed consistently well across independent datasets derived from both natural accessions and breeding populations. Although the inclusion of SVs yielded only modest overall gains in predictive power due to many causative SVs being in strong linkage with nearby SNPs, it improved the capture of variation poorly tagged by SNPs alone, as demonstrated by enhanced prediction accuracy for traits such as flesh color category.

Together, our results demonstrate how population-scale, SV-resolved pangenomics can provide both evolutionary insights and practical tools for trait-variant association and crop improvement. As watermelon breeding advances toward greater precision and efficiency, integrating pangenomic resources with genomics-assisted selection and genome editing will be essential for harnessing wild genetic diversity and accelerating the development of cultivars with improved fruit quality, disease resistance, and environmental adaptability.

## Methods

### Plant materials and phenotyping

Cultivated and wild watermelon accessions were obtained from Beijing Vegetable Research Center (BVRC) and the U.S. National Plant Germplasm System (NPGS). For phenotyping, accessions were planted in triplicate at the Hainan Experiment Station of BVRC (18° 27′ N, 108° 57′E) in 2022. One fruit per plant was harvested 34 days after pollination, with three biological replicates per accession. Each fruit was cut longitudinally, photographed, and sampled to determine flesh sugar content, flesh color intensity, rind firmness, fruit length, and fruit width. Flesh sugar content was measured at the center of the flesh in °Brix using a hand-held digital PAL-1 refractometer (Atago, Bellevue, WA, USA). Flesh color intensity was assessed from fruit images with a colorimeter (Minolta CR-400, Tokyo, Japan) to measure CIE L*, a*, b*, C* (chroma) and h* values. Rind firmness was measured at the equatorial region of each fruit using a hand-held fruit sclerometer with a 3.0 mm diameter tip (FR-5120, Lutron Electronic Enterprise Co., Ltd., Taiwan), and flesh firmness was measured at the center flesh. The fruit shape index was calculated as the ratio of fruit length to fruit width.

### Genome sequencing and assembly

High-molecular-weight genomic DNA was extracted from young fresh leaves using the cetyltrimethylammonium bromide (CTAB) method^51^. SMRTbell libraries were prepared using the SMRTbell Express Template Prep Kit 2.0 (PacBio) and sequenced on the PacBio Sequel II platform in circular consensus sequencing (ccs) mode to generate high-fidelity (HiFi) reads. For the selected ten accessions, ONT and Hi-C sequencing libraries were constructed according to the manufacturers’ instructions and sequenced on the Oxford Nanopore PromethION and Illumina NovaSeq platforms, respectively. HiFi reads for each accession were processed using HiFiAdapterFilt^52^ to remove adapter sequences. The cleaned HiFi reads, together with ONT and Hi-C data when available, were assembled into contigs using hifiasm^53^. Haplotypic duplications were removed using the purge_dups package^54^. Potential contaminant sequences from microorganisms and organelle genomes were identified and removed by comparing the contig sequences against the NCBI nt/nr database^55^. For accessions with Hi-C data, pseudochromosomes were constructed using the 3D-DNA software^56^. For the remaining accessions, chromosome-level assemblies were generated using RagTag^57^, guided by previously published reference genomes^2,6^. BUSCO completeness of the assembled genomes was estimated using the embryophyta_odb10 database^15^. Base accuracy was assessed using Merqury^16^. The presence of telomeres was determined by quarTeT^58^ and alignments to telomere-related repeat unit^59^.

### Genome annotation

Repeat sequences were predicted in each assembly using the EDTA pipeline^60^. These repeat sequences, along with previously generated custom repeat libraries for different *Citrullus* species^2^, were combined and processed to remove redundancy using the cleanup_nested.pl script from the EDTA package. The resulting non-redundant *Citrullus* repeat library was then used to mask repeats in the assembled genomes. Protein-coding genes were predicted in each genome using the MAKER pipeline^61^, incorporating evidence from transcript mapping, protein homology, and *ab initio* gene predictions. To prepare transcript evidence, *de novo* transcript assemblies were generated with Trinity^62^ for each species using RNA-seq data from various tissues obtained from NCBI SRA database (**Supplementary Table 18**). Furthermore, PacBio Iso-Seq full length cDNA sequences from our previous study^6^ and coding sequences of protein-coding genes from published watermelon genomes, were also incorporated as transcript evidence. Proteome sequences from cucumber^63^, melon^64^, pumpkin^65^, chayote^66^, snake gourd^67^, wax gourd^68^, and Arabidopsis^69^, as well as proteins from the Swiss-Prot database^70^ were used as protein homology evidence. AUGUSTUS^71^ and SNAP^72^ were used for *ab initio* gene predictions.

To further improve gene predictions across previously published and newly developed watermelon genomes, predicted genes from each genome were mapped to other genomes using Liftoff^73^. For each genome, coding sequences from all other genomes were projected onto it requiring at least 90% sequence identity and coverage. The best gene models were then selected from the original and projected sets using EVidenceModeler^74^. For functional annotation, protein sequences of predicted genes in each genome were aligned against the Swiss-Prot, TrEMBL, and TAIR10 databases using DIAMOND^75^, followed by assigning human readable functional descriptions using AHRD (https://github.com/groupschoof/AHRD). The Blast2GO suite^76^ was utilized for GO annotation and functional enrichment analysis. Putative nucleotide-binding site (NB-ARC) domain-containing genes were identified using the RGAugury pipeline^77^ (v 2.2).

### Chromosomal rearrangement and gene flow detection

Each genome assembly was aligned to the ‘97103’ reference genome using AnchorWave^78^ to identify collinear blocks. Additionally, gene-based syntenic blocks were detected using the R package GENESPACE^79^. Syntenic blocks identified by both programs were used to infer inter-chromosomal translocations and large inversions (≥ 100 kb), followed by manual inspection based on HiFi read and genome alignments.

Gene flow from *C. mucosospermus* to cultivated watermelon was detected using a composite-likelihood approach implemented in TreeMix^80^, with *C. colocynthis* used as the outgroup. Subsequently, genomic regions in cultivated watermelon that were introgressed from *C. mucosospermus* were identified using the ABBA-BABA test (D-statistic), as previously described^31^. Specifically, non-overlapping 100-kb windows across the genome with the top 5% *f_d_* values (degree of introgression) were defined as introgressed regions.

### Gene-based super-pangenome construction and phylogenetic analysis

Gene families across the 138 high-quality *Citrullus* genomes were identified using OrthoFinder^81^ (v2.5.5). Synteny information was then used to separate paralogous genes that were not located within the same syntenic regions. Syntenic gene blocks between each pair of genomes were detected using MCScanX^82^, and the synonymous substitution rates (*Ks*) between syntenic genes were calculated using the Yang-Nielsen algorithm implemented in the PAML package^83^. Syntenic orthologous gene blocks were used to refine the OrthoFinder-defined orthologous groups, dividing them into syntenic orthologous gene families and singletons. Based on their presence across the 138 genomes, syntenic orthologous gene families were classified into four categories: core (present in all 138 genomes), softcore (present in 137 genomes), shell (present in 2-136 genomes), and private (present in only one genome).

To reconstruct the species tree, single-copy orthologous genes (SCOs) were identified by clustering predicted proteins from 13 species, including seven *Citrullus* species, as well as bottle gourd^84^, cucumber^63^, melon^64^, snake gourd^67^, bitter gourd^85^, and walnut^86^. Protein-guided multiple coding sequence alignments of SCOs were obtained using TranslatorX^87^. Divergence times among species were estimated using BEAST2 with the ‘Optimised Relaxed Clock’ model, calibrated with known divergence times for Fagales-Cucurbitales (85.6-109 million years ago [Mya]), *Cucumis*-*Citrullus* (16.4-24.2 Mya), and cucumber-melon (5.96-13.1 Mya)^88^. The time of domestication, represented by the divergence between *C. lanatus* subsp. *cordophanus* and cultivated watermelon, was estimated using the SMC++ program^89^.

### Genome resequencing of watermelon core accessions

A watermelon core collection comprising 323 representative wild and domesticated accessions was constructed from ~1,400 accessions using GenoCore^90^, based on SNPs derived from our previously reported GBS data^91^, eight of which were also included in *de novo* genome assemblies. Additionally, accessions harboring important breeding traits were also included in the core collection. Genomic DNA was extracted from young leaf samples of the core accessions using the Qiagen DNeasy Plant Kit. Shotgun DNA libraries were constructed from the extracted DNA and sequenced on the BGISEQ-500 platform to generate 150-bp paired-end reads. Genomic resequencing data from an additional 463 watermelon accessions were retrieved from previous studies^2,6,91^. Raw sequencing reads were processed to remove adapter sequences and low-quality bases using Trimmomatic (v0.38)^92^.

### Graph pangenome construction and SV genotyping

Each of the 137 genomes was aligned to the ‘97103’ reference genome using AnchorWave^78^. Structural variants (SVs), including large insertions, deletions, and inversions, as well as SNPs and small indels, were identified based on the alignments using NucDiff^93^. Deletions longer than 10 kb were further validated based on HiFi read coverage. SVs identified from the 137 accessions were merged using bcftools^94^ (v1.14), and redundant variants were removed using the script findDup.R (https://github.com/vgteam/giraffe-sv-paper/tree/master/scripts/sv/remap-to-dedup-merged-svs). A pangenome graph was constructed using PanGenie^95^ (v3.0.1) from the identified SVs, SNPs, and small indels, with the ‘97103’ genome used as the reference. SVs in the graph pangenome were then genotyped in the resequenced accessions using PanGenie with the cleaned resequencing reads.

### Genome-wide association studies

SNPs in all resequenced *Citrullus* accessions were called using the Sentieon package (https://www.sentieon.com/), followed by hard filtering with recommended parameters^96^. A total of 20,210,332 bi-allelic SNPs with both missing rates and heterozygous rates below 20% were retained. These SNPs were then combined with all identified SVs from the graph pangenome and used for GWAS. Five accessions from *C. naudinianus*, *C. rehmii*, and *C. ecirrhosus* were excluded because of small sample sizes and lack of phenotypic data. For each trait, variants with a high missing rate (>30%) or low minor allele counts (<10) among accessions with phenotypic data were removed. To account for population structure, a kinship matrix was generated using FaST-LMM^97^ (v2.07), and GWAS was performed using the linear mixed model implemented in FaST-LMM. Genome-wide significance thresholds were determined by calculating the effective number of independent variants using the Genetic type 1 Error Calculator^98^ (GEC v0.2).

### Genomic prediction

To generate marker panels for genomic prediction of each trait, we applied CropGBM^43^ (v1.1.2) to perform feature selection using SNP-only and SNP+SV variant sets. Prior to feature selection, variants with a minor allele frequency (MAF) < 0.05 or missing rate greater than 20% were excluded to retain high-confidence variants. Additionally, linkage disequilibrium (LD) pruning was performed to remove highly correlated variants, using an *r*^2^ threshold of 0.999. The resulting marker panels were used to generate genomic prediction models with CropGBM. Five-fold cross-validation was used with five repeats to evaluate model performance.

### Genetic mapping of flesh color intensity trait

To map loci controlling flesh color intensity, two independent F_2_ populations were developed from crosses of cultivars ‘Ming 58’ (scarlet red flesh) × ‘JX2’ (pink flesh) and ‘JLM’ (yellow flesh) × ‘Cream of Saskatchewan’ (pale yellow flesh; hereafter ‘CS’). Flesh color intensity was determined at 34 days after pollination (DAP). For bulk segregant analysis (BSA), two DNA pools were constructed from the ‘Ming 58’ and ‘JX2’ F_2_ population: one comprising 20 individuals with the lowest chroma values and the other comprising 20 individuals with the highest chroma values. Genomic DNA was extracted from each individual using the CTAB method, mixed in equal amount within each pool, and subjected to library construction and whole-genome sequencing on the Illumina HiSeq platform. Variant calling was performed using the Sentieon package (https://www.sentieon.com/), followed by hard filtering with recommended parameters^96^. BSA was performed using the R package QTLseqr^99^ to identify QTLs of flesh color intensity. To perform fine mapping of the candidate region, larger F_2_ segregating populations were generated, consisting of 636 individuals from the cross of ‘JLM’ × ‘CS’ and 141 individuals from the cross of ‘Ming 58’ × ‘JX2’. Based on SNPs identified between parental lines, Kompetitive Allele Specific PCR (KASP) markers (**Supplementary Table 19**) were developed. The linkage map was constructed from KASP markers in each population using QTL IciMapping^100^ (v4.2).

### Quantitative RT-PCR and transient dual-luciferase activity assay

Total RNA was extracted using the Quick RNA isolation kit (Huayueyang Biotechnologies Co., Ltd.). First-strand cDNA was synthesized from 1 ug of total RNA using SuperScript^TM^ III Reverse Transcriptase (Invitrogen, Carlsbad, CA, USA) with oligo(dT)18 primers, according to the manufacturer’s instructions. Gene expression levels were quantified using SYBR Green-based qPCR on a Roche LightCycler® 480 system (Roche, Basel, Switzerland). Three biological replicates were performed for each gene, with watermelon *ClActin1* gene used as the internal reference. Relative expression was calculated using the 2^−ΔΔCt^ method after normalization against *ClActin1*.

To compare the promoter activity associated with different copy numbers of the 1,258-bp sequence upstream of *ClFCI1*, one, two, and three copies of this sequence were PCR-amplified from ‘CS’ and ‘JLM’ using primers FCI-SV-PstI/BamHI (**Supplementary Table 20**). The resulting PCR fragments were cloned into the PstI and BamHI sites of the pGreenII 0800-LUC vector. The constructs were then transformed into watermelon fruit protoplasts following the method described previously^101^. Luciferase activity was measured using the Dual-Luciferase Reporter Assay Kit following the manufacturer’s instructions (Vazyme Biotech, China). Ten biological replicates were performed for each construct.

### *Agrobacterium*-mediated transformation

The full-length cDNA and a partial cDNA fragment of *ClFCI1* were amplified using FCI-PacI/AscI and FCI-AscI/PacI primers, respectively (**Supplementary Table 20**). These PCR products were cloned into the PacI/AscI sites of pMDC85 (ref. ^102^) to generate *ClFCI1* overexpression and knockdown constructs, respectively. The resulting constructs were then introduced into *Agrobacterium tumefaciens* strain C58/ATCC 33970. Plant transformation was performed as previously described^101^. Transgene insertion in the transformed watermelon lines was confirmed by PCR using the AS013 PAT/bar Kit (Envirologix Inc., Portland, ME, USA).

### Characterization of *ClFCI1* transgenic lines

Carotenoids were extracted from mature fruit flesh (5 g; 34 DAP) of *ClFCI1* overexpression and knockdown lines using a hexane:acetone:ethanol (50:25:25, v/v/v) mixture. Carotenoid composition and content were then determined using a Nexera HPLC system (Shimadzu).

For transcriptome analyses, total RNA was extracted from the fruit flesh of *ClFCI1* knockdown and overexpression lines, as well as the wild-type line (‘ZZJM’). RNA-Seq libraries were constructed from total RNA using the TruSeq^TM^ RNA Sample Prep Kit (Illumina, USA) and sequenced on the Illumina HiSeq 4000 platform to generate paired-end 150-bp reads. Three biological replicates were conducted for each sample. Raw RNA-Seq reads were cleaned using Trimmomatic^92^, and the cleaned reads were aligned to the ‘97103’ reference genome using HISAT2 (ref. ^103^). Raw counts for each protein-coding gene were calculated using featureCounts^104^ and then normalized to transcripts per million (TPM). Differentially expressed genes (DEGs) were identified using DESeq2 (ref. ^105^) by comparing *ClFCI1*-knockdown and *ClFCI1*-overexpression lines to wild-type fruits. The Benjamini-Hochberg method^106^ was used to control the false discovery rate (FDR). Genes with FDR < 0.05 and |log₂(fold change)| ≥ 1 were considered significantly differentially expressed.

## Data availability

Raw HiFi, ONT, Hi-C, and genome resequencing reads have been deposited in the NCBI BioProject database under accession number PRJNA1272048. Genome assemblies and annotations, and variant files in VCF format are available at CuGenDBv2 (http://cucurbitgenomics.org/v2/ftp/pan-genome/watermelon/graph_pangenome/).

## Author contributions

Z.F. and Y.X. designed and supervised the project. S.G., S.A.H., C.M., R.J., S.E.B., P.W., C.K., A.L. and R.G. contributed to sample collection and DNA extraction. S.G., H.S., S.Liao, J.Zhang, R.J. and Z.F. coordinated genome sequencing. S.G., S.Liao, J.Zhang, G.G., J.W., Y.Y., Y.R., S.T., S.Li and H.Z. performed phenotyping for fruit-quality traits. H.S., Z.Z., X.Z. and S.W. contributed to genome assembly and annotation, as well as pangenome and population genetic analyses. H.S. and J.Zhao conducted the genomic prediction analysis. J.Zhang, H.S. and S.Liao contributed to genetic mapping and gene functional characterization. H.S., Z.Z., J.Zhang., X.Z, and S.W. wrote the manuscript. Z.F. and Y.X. revised the manuscript.

## Acknowledgements

We thank Susanne S. Renner for providing seeds of *C. lanatus* subsp. *cordophanus*. This research was supported by grants from Beijing Rural Revitalization Agricultural Science and Technology Project (NY2401130025), National Natural Science Foundation of China (Grant No. 32172592, 32330093), the Scientific and Technological Innovation Capacity Building Project of BAAFS (KJCX20251008), the Scientist Training Program of BAAFS (JKZX202401), Ministry of Agriculture of China (CARS-25), USDA National Institute of Food and Agriculture Specialty Crop Research Initiative (2015-51181-24285 and 2020-51181-32139).

