## Supplementary Figures for "Population-level super-pangenome reveals genome evolution and empowers precision breeding in watermelon"

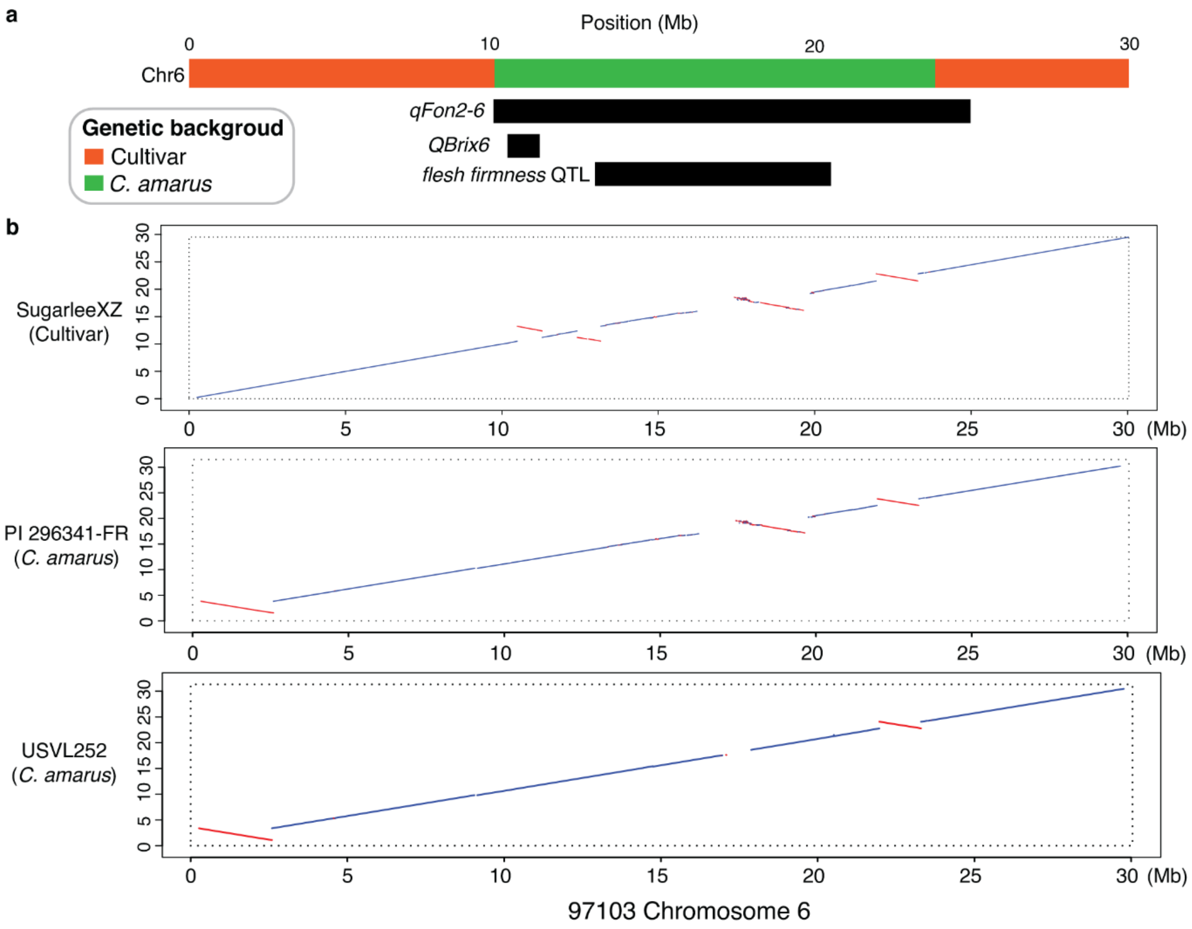


**Supplementary Fig. 1. Large inversions associated with recombination suppression.** (**a**) A genomic region on chromosome 6 (green box) in cultivated watermelon introgressed from *C. amarus*, overlapping with three QTLs related to Fusarium wilt resistance (*qFon2-6*), flesh sweetness (*QBrix6*), and flesh firmness. (**b**) Alignments of chromosome 6 between reference '97103' and the accessions 'SugarleeXZ', 'PI 264341-FR', and 'USVL252', all of which harbor resistance to Fusarium wilt.

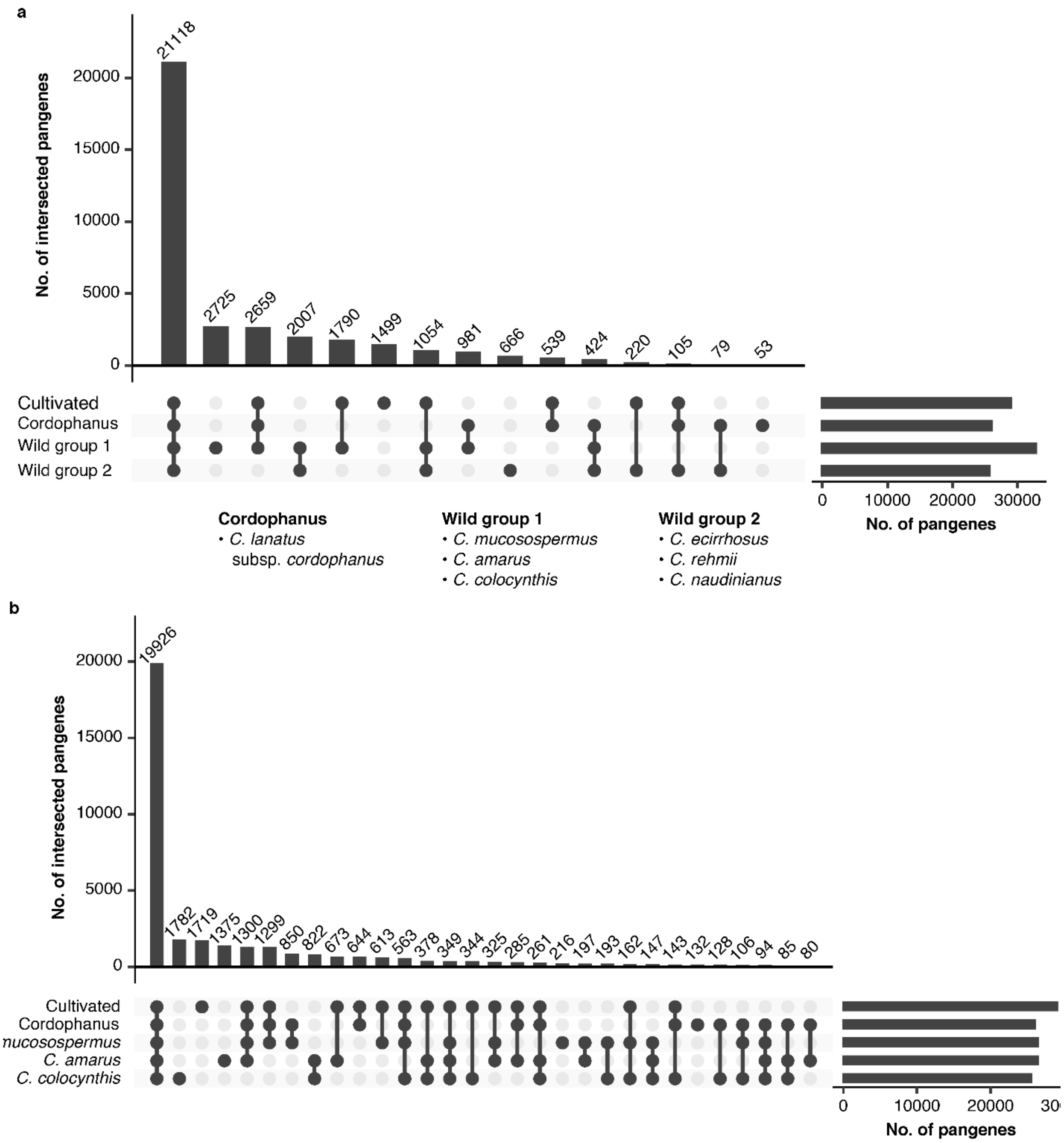


**Supplementary Fig. 2. Pangene distribution across *Citrullus* groups.** (**a**) UpSet plot showing intersections of pangenes among different *Citrullus* species and groups. (**b**) UpSet plot showing intersections of pangenes among cultivated watermelon, its direct progenitor (*C. lanatus* subsp. *cordophanus*), and three wild species commonly used in watermelon breeding.

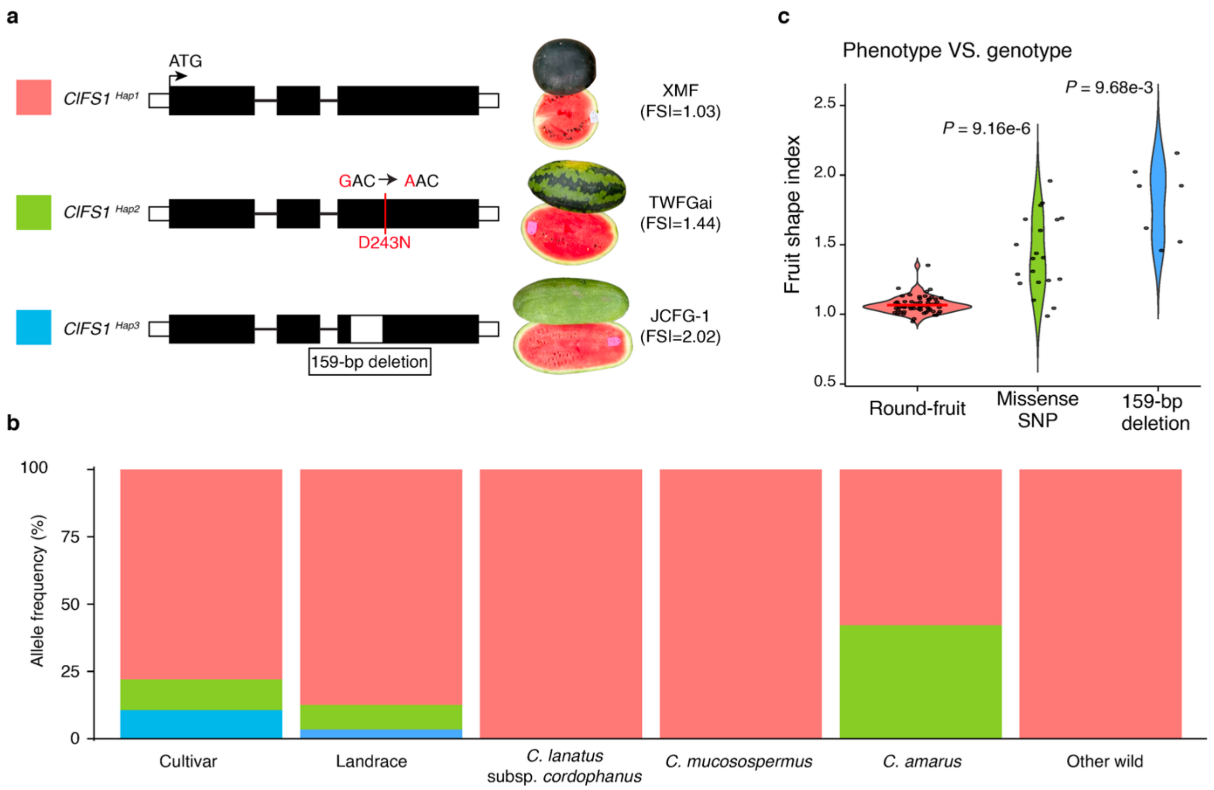


**Supplementary Fig. 3. Haplotypes of the fruit shape-related gene *ClFS1*.** (**a**) Structure diagram of the missense SNP (*ClFS1*$^{Hap2}$) and 159-bp deletion (*ClFS1*$^{Hap3}$) in *ClFS1*. Representative fruits carrying the corresponding haplotypes are shown. *ClFS1*$^{Hap1}$ denotes the reference allele from '97103'. (**b**) Allele frequencies of the three haplotypes across *Citrullus* groups. (**c**) Effect of different haplotypes of *ClFS1* on fruit shape.

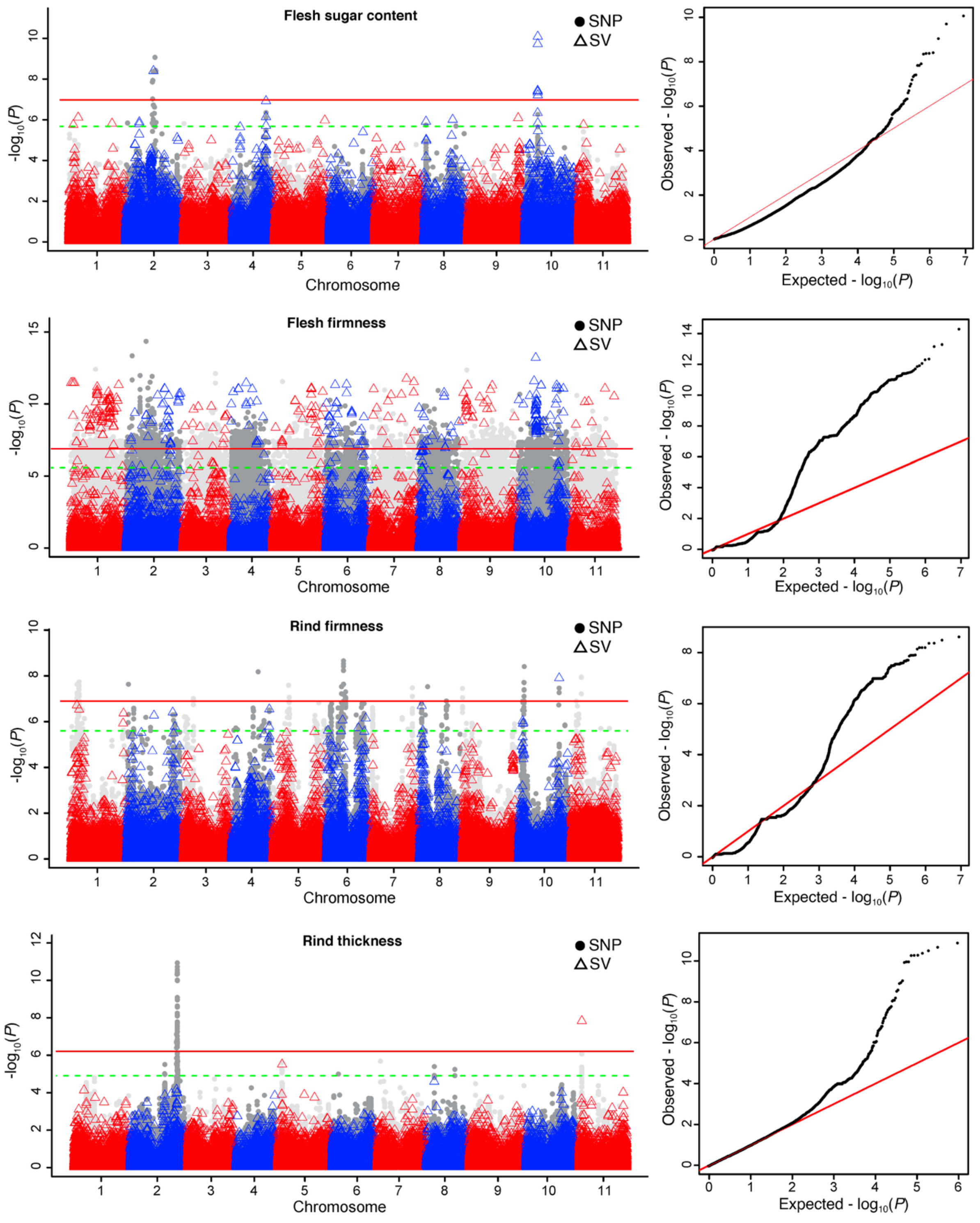


**Supplementary Fig. 4. Genome-wide association studies (GWAS) of watermelon fruit quality and disease resistance traits**. Manhattan plots (left) and quantile-quantile (Q-Q) plots (right) of GWAS are shown for each trait. Red and green horizontal lines indicate genome-wide significance thresholds at α = 0.05 and α = 0.10, respectively.

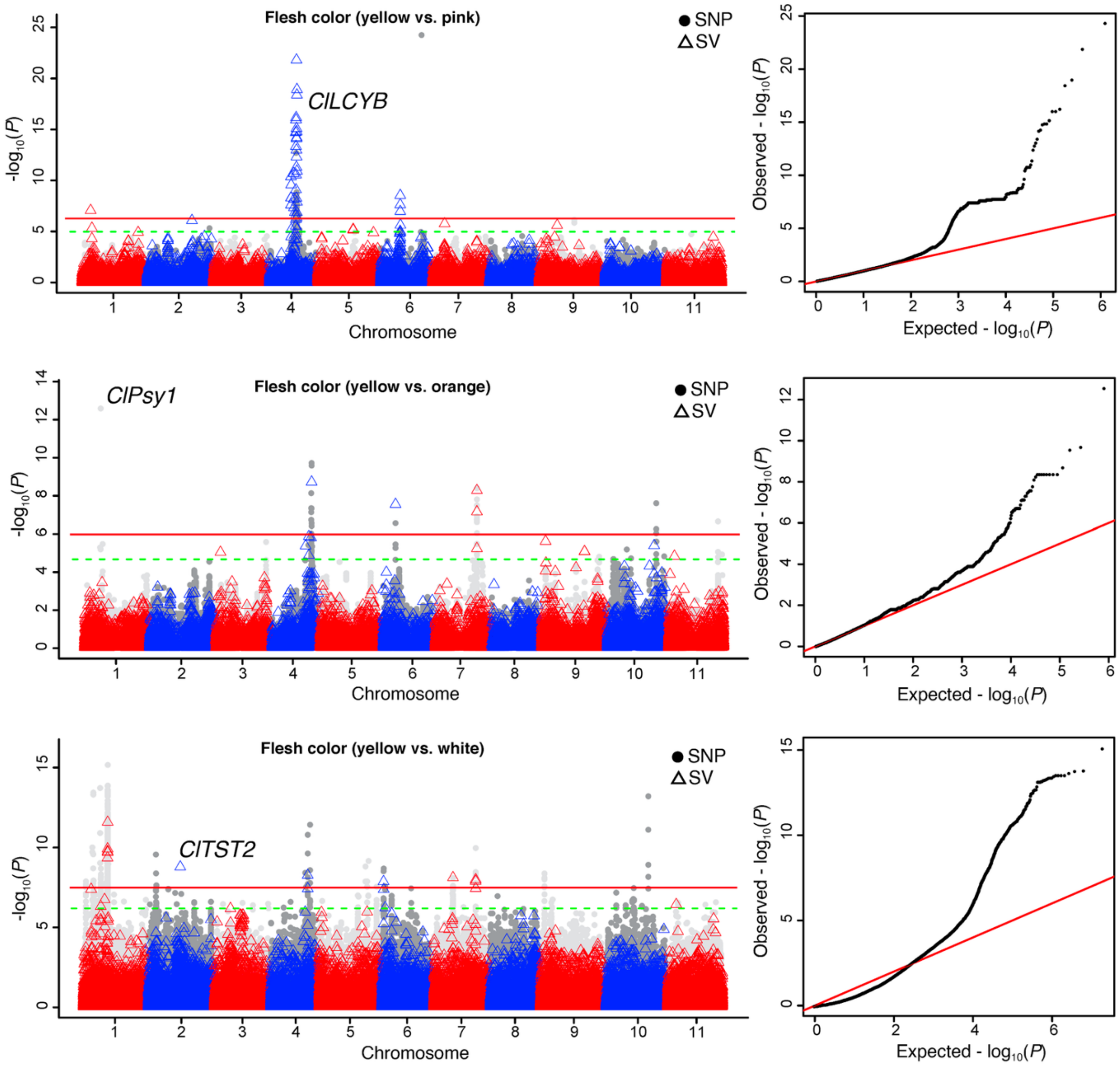


**Supplementary Fig. 4. Continued**

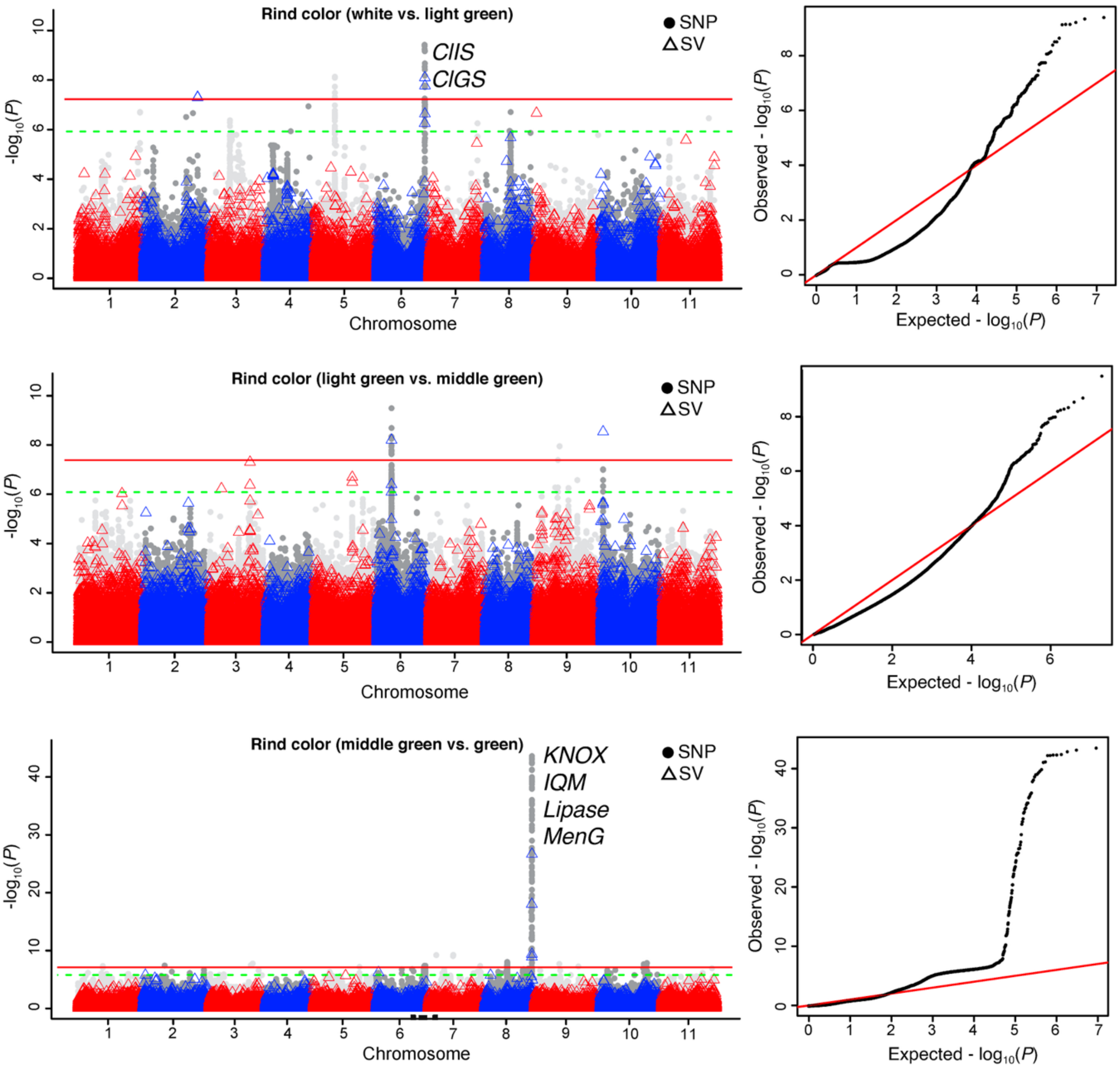


**Supplementary Fig. 4. Continued**

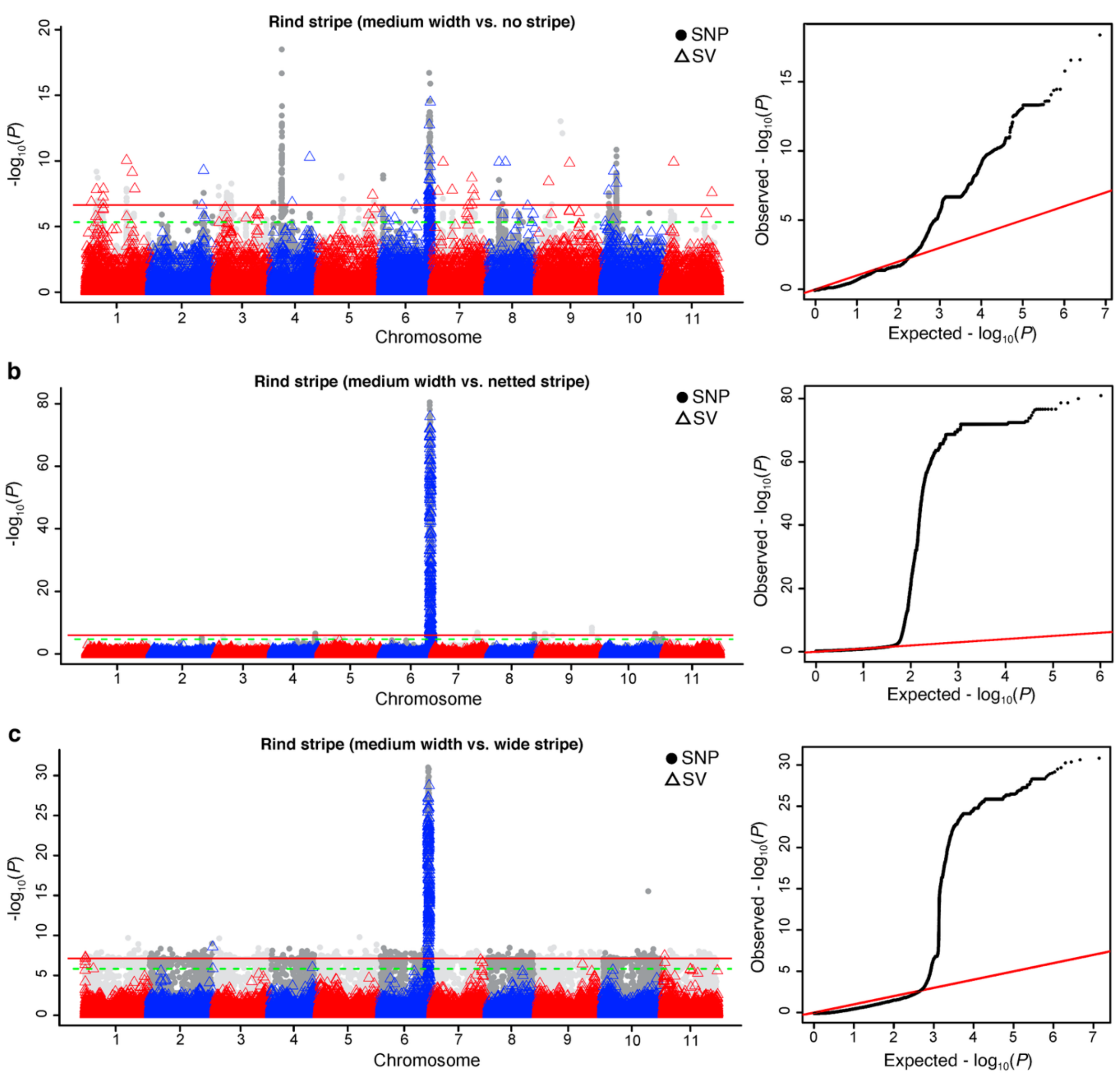


**Supplementary Fig. 4. Continued**

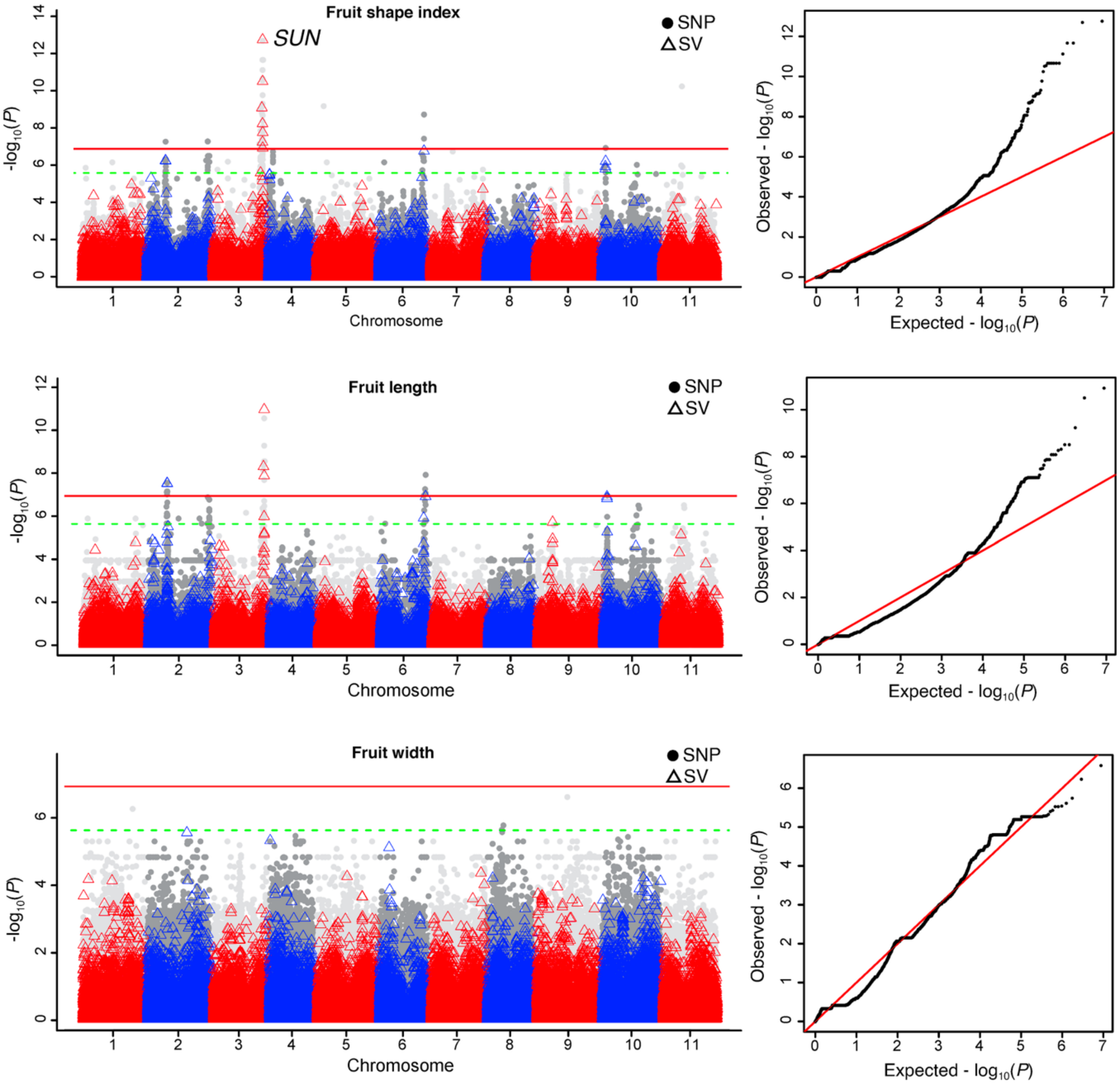


**Supplementary Fig. 4. Continued**

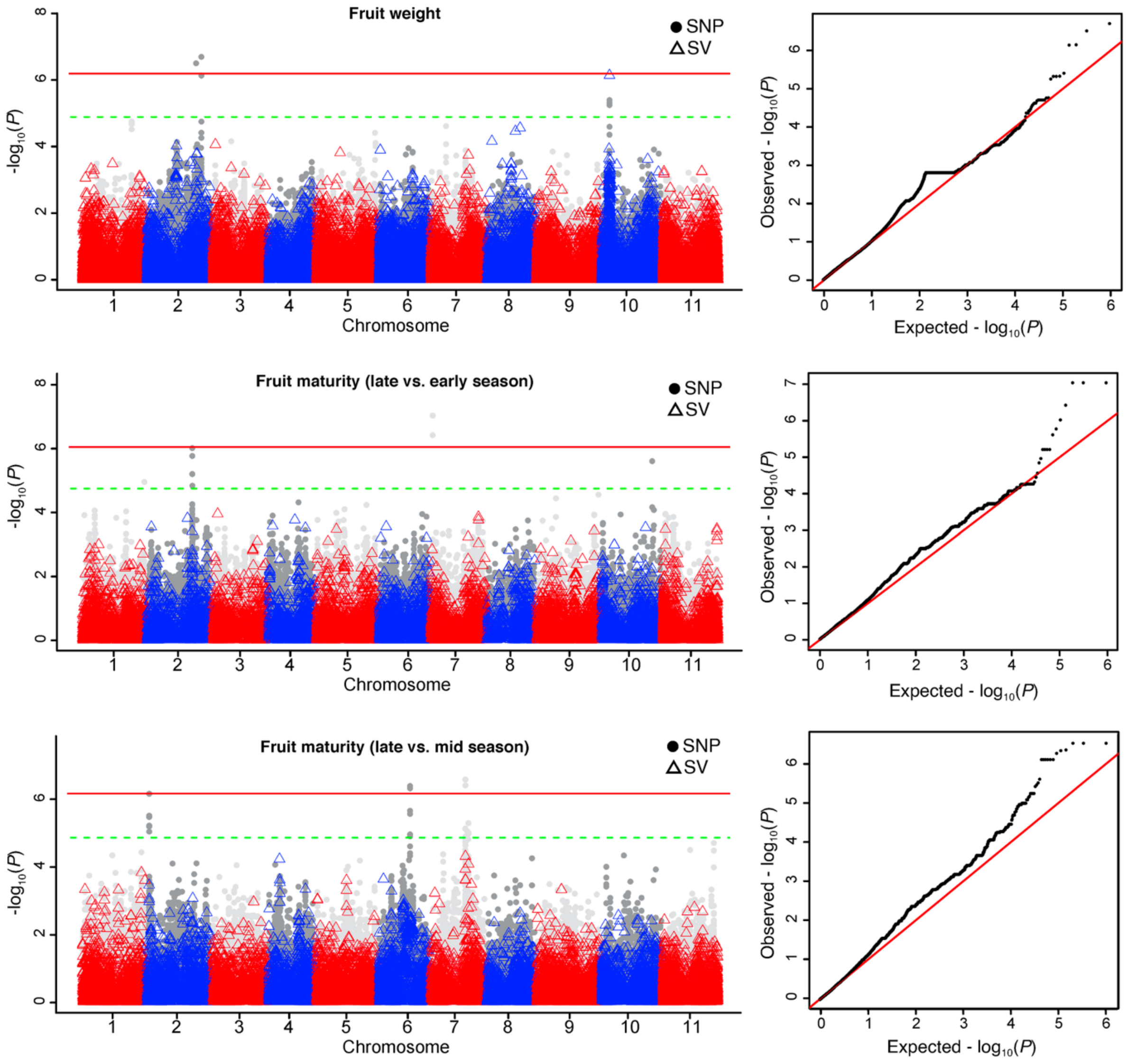


**Supplementary Fig. 4. Continued**

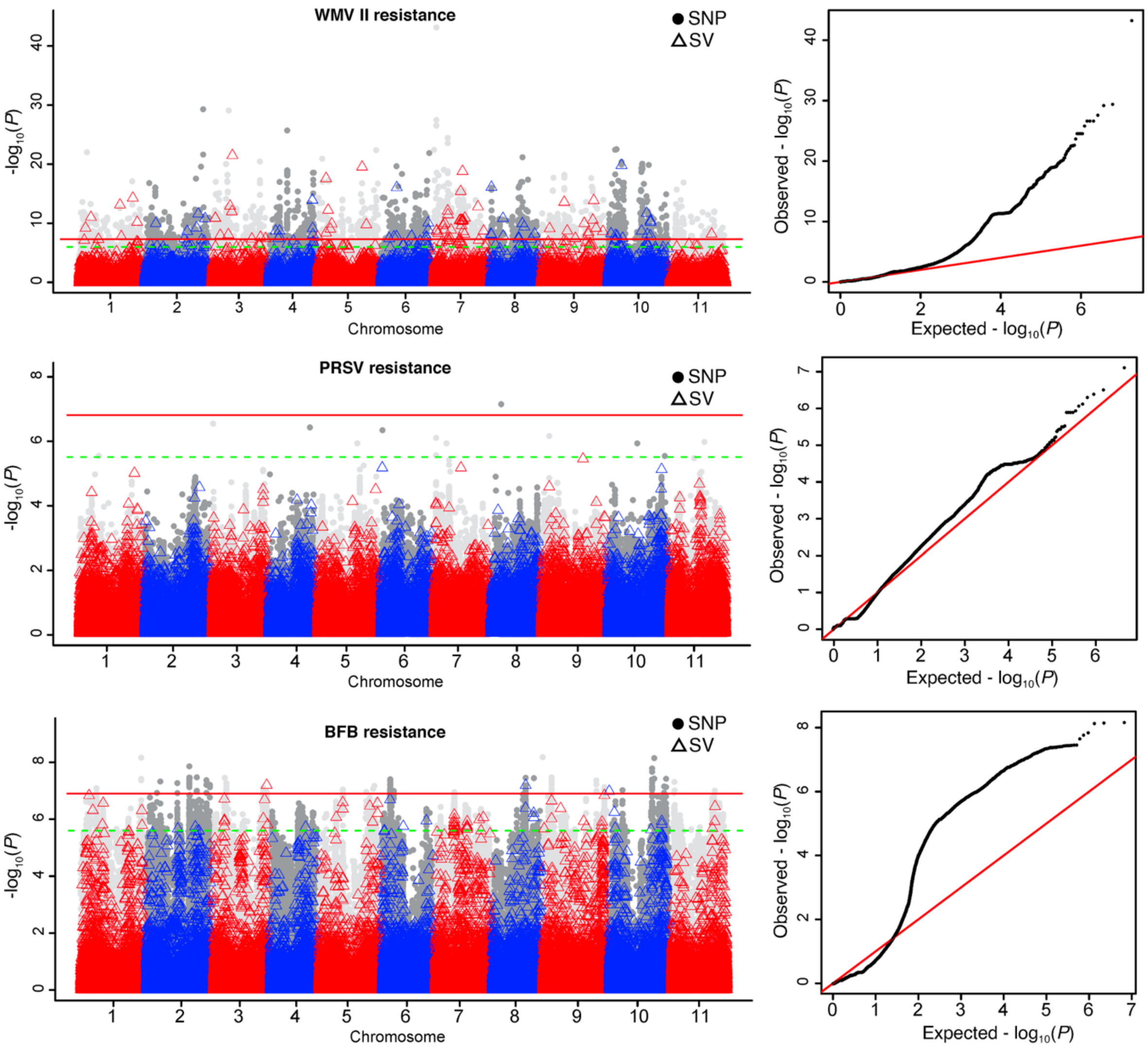


**Supplementary Fig. 4. Continued**

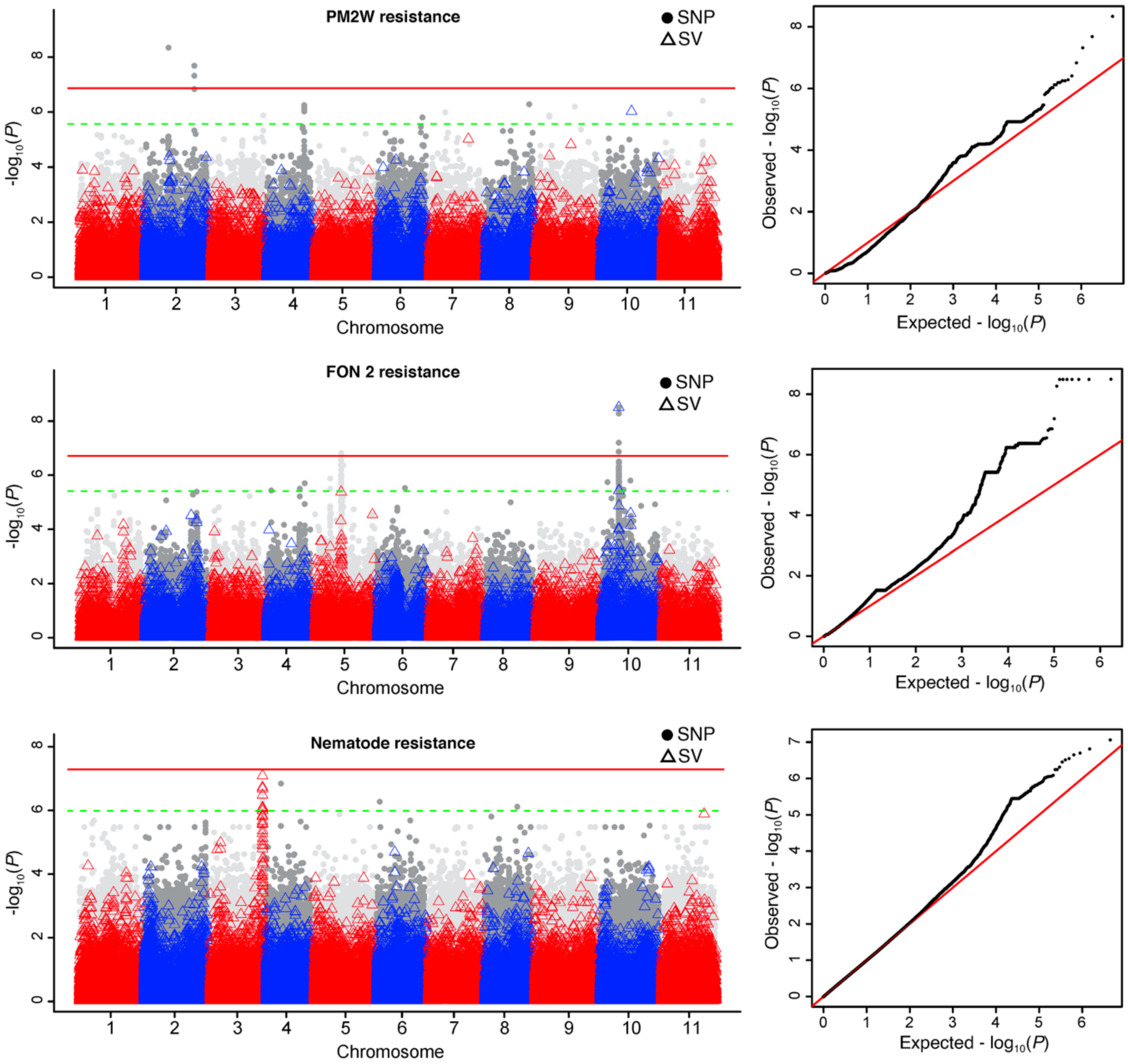


**Supplementary Fig. 4. Continued**

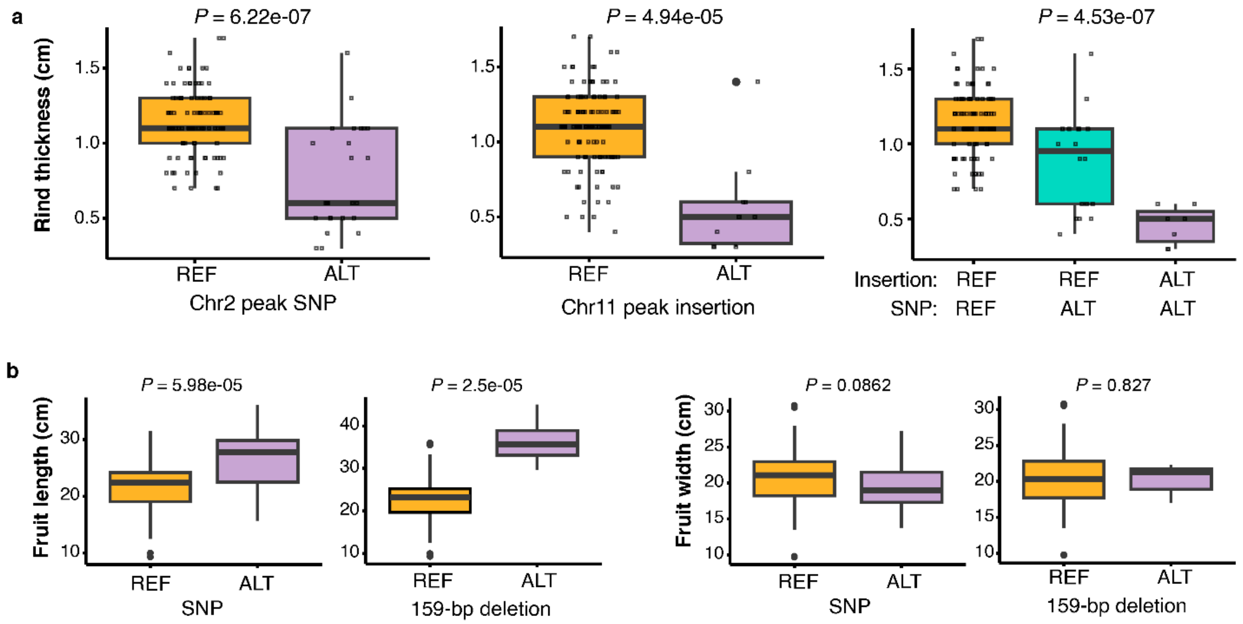


**Supplementary Fig. 5.** (**a**) Box plots of rind thickness in accessions carrying the reference (REF) and alternative (ALT) alleles at the peak SNP (left) on chromosome 2 and the peak 20-bp insertion (INS) (middle) on chromosome 11, as well as their combinations (right). (**b**) Box plots of fruit length and fruit width in accessions carrying the REF and ALT alleles at the missense SNP (left) and the 159-bp deletion (right) in the *SUN* gene. P values were calculated using the Kruskal-Wallis tests. For each boxplot, the lower and upper bounds indicate the first and third quartiles, respectively, the center line indicates the median, and the whiskers extend to 1.5× the interquartile range.

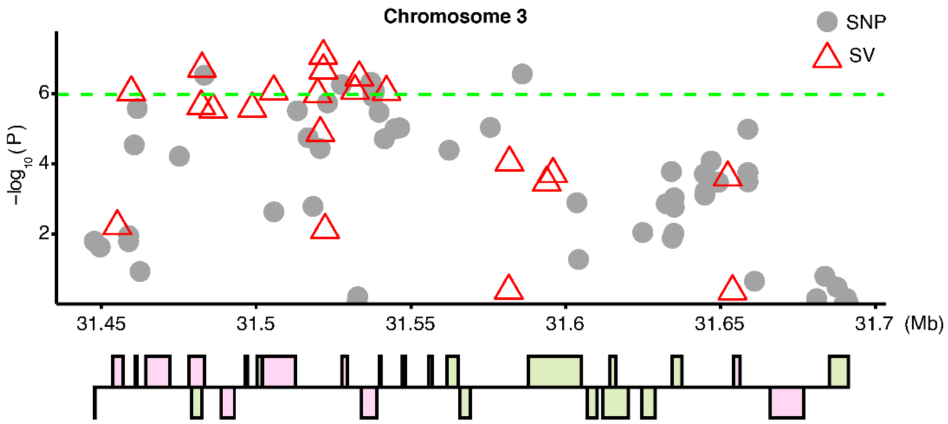


**Supplementary Fig. 6. Nematode resistance-associated SVs in the *CBP60* gene cluster.** Top: Manhattan plot of GWAS for nematode resistance using SNP (gray point) and SV (red triangle) variants. Bottom: Genes in the genomic region associated with nematode resistance. *CBP60* family genes are shown in green and other genes are shown in pink.

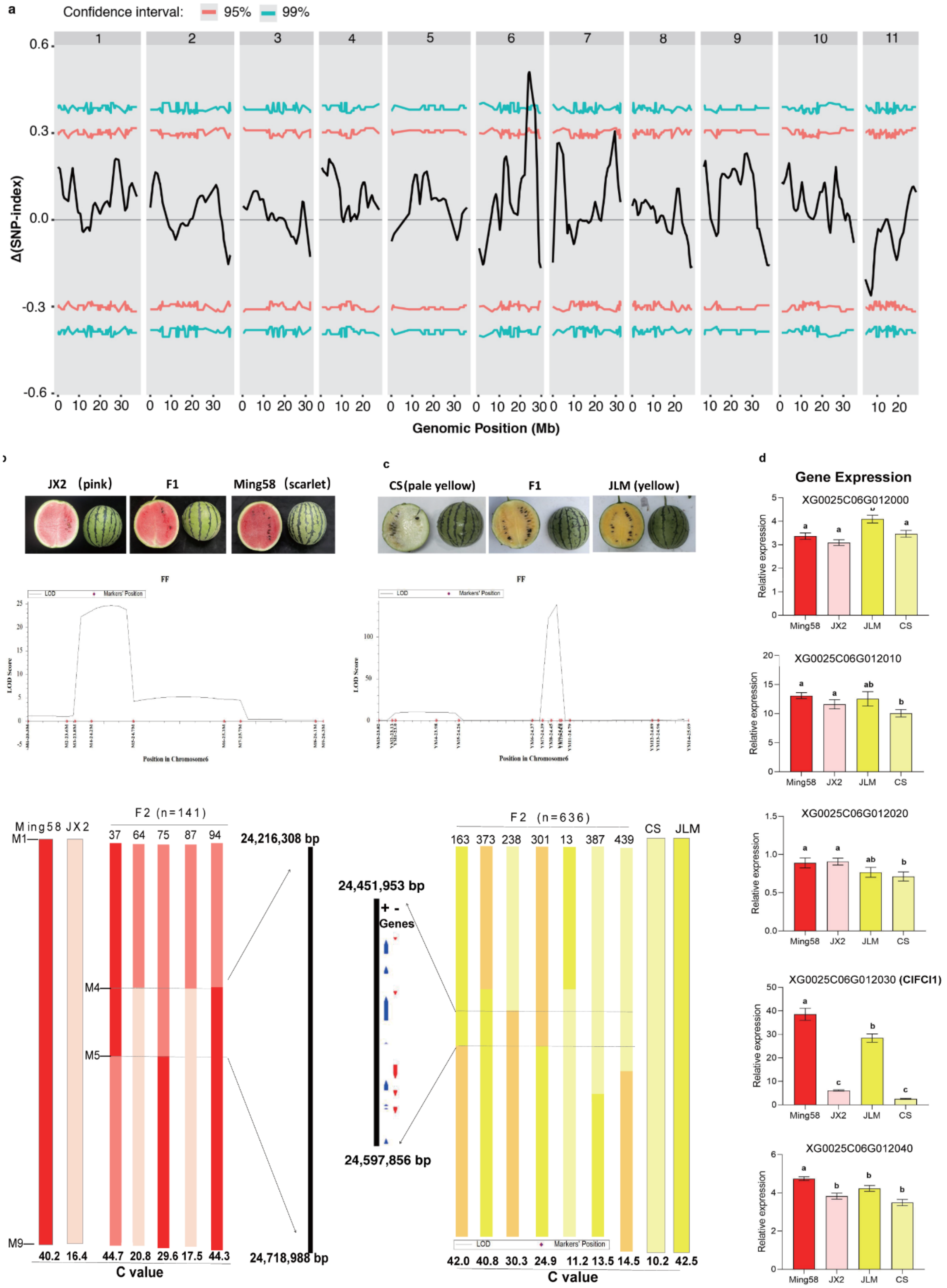


**Supplementary Fig. 7. Genetic mapping of flesh color intensity in watermelon. (a)** Genome-wide ΔSNP-index profile from bulked-segregant analysis (BSA) of an F2 population derived from the cross ‘Ming 58’ (scarlet flesh) × ‘JX2’ (pink flesh). The black curve represents ΔSNP-index values, while red and blue envelopes mark the 95% and 99% confidence thresholds, respectively. **(b,c)** Recombinant-based fine-mapping of the *ClFCI1* locus in 141 F2 plants from the 'Ming 58' × 'JX2' cross (**b**), and 636 F2 plants from the 'JLM' × 'Cream of Saskatchewan' (CS) cross (**c**). Top**,** fruits of parental lines and F1 hybrids. Middle**,** QTL curves showing logarithm of odds (LOD) scores using KASP markers between 23 Mb and 26 Mb on chromosome 6. Bottom**,** graphical genotypes of key recombinant individuals. Pink, red, and scarlet segments denote homozygous 'JX2', heterozygous, and homozygous 'Ming 58' regions, respectively; pale yellow, orange, and yellow segments denote homozygous 'CS', heterozygous, and homozygous 'JLM' regions, respectively. **(d)** Relative expression levels of candidate genes in the four parental lines ('Ming 58', 'JX2', 'JLM', and 'CS').
